# A Single-Cell Transcriptome Atlas for Zebrafish Development

**DOI:** 10.1101/738344

**Authors:** Dylan R. Farnsworth, Lauren Saunders, Adam C. Miller

## Abstract

The ability to define cell types and how they change during organogenesis is central to our understanding of animal development and human disease. Despite the crucial nature of this knowledge, we have yet to fully characterize all distinct cell types and the gene expression differences that generate cell types during development. To address this knowledge gap, we produced an Atlas using single-cell RNA-sequencing methods to investigate gene expression from the pharyngula to early larval stages in developing zebrafish. Our single-cell transcriptome Atlas encompasses transcriptional profiles from 44,102 cells across four days of development using duplicate experiments that confirmed high reproducibility. We annotated 220 identified clusters and highlighted several strategies for interrogating changes in gene expression associated with the development of zebrafish embryos at single-cell resolution. Furthermore, we highlight the power of this analysis to assign new cell-type or developmental stage-specific expression information to many genes, including those that are currently known only by sequence and/or that lack expression information altogether. The resulting Atlas is a resource of biologists to generate hypotheses for genetic (mutant) or functional analysis, to launch an effort to define the diversity of cell-types during zebrafish organogenesis, and to examine the transcriptional profiles that produce each cell type over developmental time.

## INTRODUCTION

During animal development, stem cells generate a vast diversity of differentiated cells to form functioning tissues and organs. The specification of cell fates during development and the ability to function properly during adult life are determined by the RNA messages cells express and the functions and quantities of proteins produced from these messages. A central question in biology is how gene expression unfolds over time to precisely coordinate the development of adult organisms. Such coordinated gene expression programs are critical given that disruptions to the expression of cell-type specific RNAs can lead to abnormal function and the progression of degeneration and disease. A major obstacle for regenerative therapies is not knowing the full system of genes normally expressed in a given cell type, thus preventing the development of cell-type specific pharmaceuticals as well as the efficient reprogramming of stem cells to regenerate damaged tissue. Furthermore, genes associated with genetic disorders are often poorly described in terms of cell and developmental stage. To understand development, and to provide a foundation for inventing therapeutics, we must comprehend the systems-level phenomenon of how genes and proteins are used across cell type and over time.

Characterizing the *in toto* systems of a developing animal is a challenge. However, recent advances in single-cell RNA-seq (scRNAseq) provide a platform to track transcriptional changes across thousands of cells simultaneously. Capturing all transcriptional changes across all of the cells of a developing animal is a powerful step towards identifying the molecular and genetic basis of cell-type specification, organogenesis, and adult homeostasis. To initiate a collective effort to compile these data, to transcriptionally identify cell types, and afford researchers the opportunity to efficiently identify *de novo* candidates for genetic analysis in tissues, cell types, and developmental gene expression programs of interest, we have drafted an Atlas from whole zebrafish embryos and larvae during organogenesis, where single-cell transcriptomic data are compiled from samples that span four days of development (1-5 day post fertilization).

Zebrafish is an excellent model system for producing an Atlas of vertebrate gene expression over developmental time. Zebrafish development is rapid, with free-swimming animals emerging within the first three days after fertilization, accompanied by a system-wide expansion and differentiation of cell types including: neurogenesis (and associated complex behaviors), maturation of blood, muscle, gastro-intestinal tissues, germ cells, and the specification of pigmented skin epithelium, vasculature, cartilage, and bone. Here, we present an Atlas and demonstrate how to use it for insight into several of these developmental processes. We identify hundreds of transcriptionally-defined cell types and their corresponding gene expression developmental trajectories, and we anchor them to zebrafish anatomy by comparison with RNA *in situ* expression patterns. This Atlas provides a rich resource of transcriptionally-defined cell types across zebrafish organogenesis. Researchers can mine this resource for transcripts that were previously not attributed to specific cell types of interest (*de novo* gene expression analysis) as well probe for temporal changes in gene expression that underly cell-type specification during development.

## RESULTS

### The zebrafish scRNA-seq Atlas across organogenesis

We designed the Atlas to be a tool for investigating changes in gene expression associated with cell-fate specification during animal development with the goal of providing new insight into vertebrate organogenesis. To generate high-quality single-cell transcriptional profiles, we examined the reproducibility of our cell dissociation and scRNAseq methods by performing replicate dissociations derived from independent matings, cDNA library preparations, and sequencing for each stage of development profiled in this study (1, 2, and 5 days post fertilization (dpf)). These data combined to total 44,012 cells, 74,914 mean reads per cell, 1807 median genes per cell, and a median of 8345 unique transcript molecules per cell (Figure S1A,B). We used Seurat (Butler et al., 2018) to cluster transcriptionally similar cells together and compared the results of replicate experiments to each other in terms of cluster diversity and the representation of cells from each duplicate within all clusters. We found that each independent experiment contributed to the computationally identified clusters at each age, showing that our dissociation methods were reproducibly sampling across all accessible cell types (Figure S1C-E). Thus, our methodology supports a robust and reproducible single-cell transcriptomic analysis and clustering, enabling investigation of cell type, state, and developmental transitions through organogenesis

### Cell-type annotation and de novo gene identification for clusters within the Atlas

The Zebrafish scRNAseq Atlas should offer the ability to produce accurate gene expression profiles for cell types of interest at multiple developmental stages. We therefore aggregated the results of all six experiments from three distinct developmental stages (1, 2, and 5 dpf) to examine the diversity of transcriptional cell types across developmental time (Figure 1A). Aggregation resulted in 220 computationally identified clusters (Figure S2) and for each cluster we sought to assign it to the most likely cell type.

**Figure 1.**
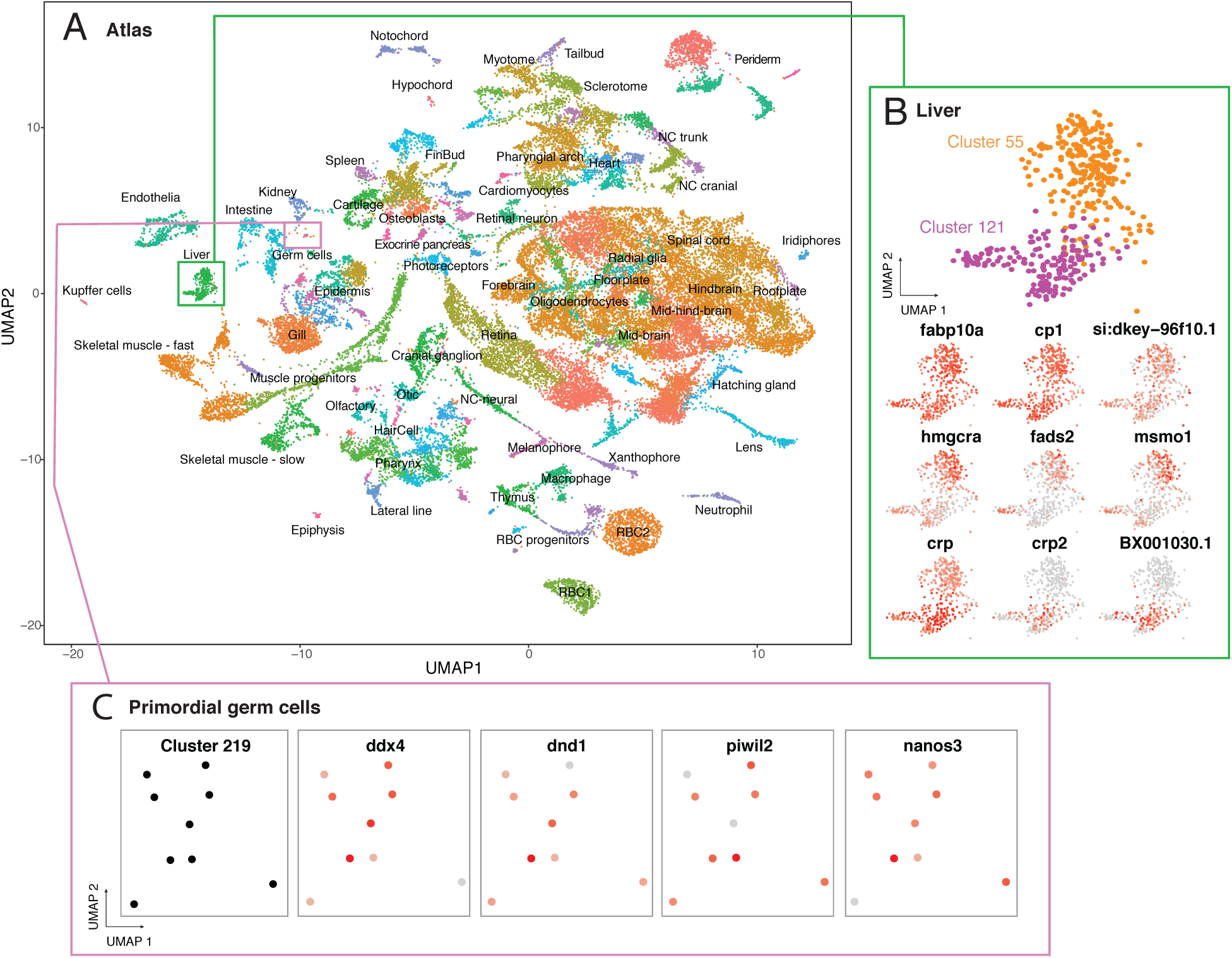
scRNA-seq Atlas of developing zebrafish embryos during organogenesis. (A) Clustering of cell types enables gene expression analysis across transcriptional cells types over developmental time. Colors correspond to labels which indicate a grouping of clusters and annotations. Green box describes clusters enlarged in B; pink box describes cluster enlarged in C. Dashed oval approximates the region described in Figure 2. (B) Heterogeneity within hepatocytes is revealed by the identification of two clusters (55 and 121) with different but related gene expression profiles. Common and differential gene expression between these clusters are plotted using red to indicate high levels of expression and grey for low expression. (C) Rare cells types, including primordial germ cells (PGCs), can be efficiently profiled and are restricted to a single cluster (219). Four markers of PGCs show high levels of expression within this cluster.

To establish a logic for annotating clusters to cell types, we first sought to use the expression of unambiguous, previously identified marker genes. For example, genes associated with hepatocytes, *fabp10a* and *cp* (Her et al., 2003; Korzh et al., 2001), were among the most differentially expressed in our entire data set, as demonstrated by their enrichment in the first principal component (PC) of the PC analysis (PCA) used to cluster cells (Figure S3). We plotted expression of these genes within the Atlas and identified two clusters in close proximity that also expressed additional transcripts expressed in and associated with liver development and function (Figure 1B, Table S1 see clusters 55 and 121). Examining the top 30 most differentially expressed genes in these clusters, we found that 15 that have been described to be expressed in the hepatocytes (The Zebrafish Information Network, ZFIN)(Howe et al., 2013), strongly supporting the annotation of these clusters (Figure 1B, Table 2, see clusters 55 and 121). Additionally, we find 7 genes that are poorly characterized, including *si:dkey-96f10.1* (Fig. 1B), that are expressed in both hepatocyte clusters, highlighting the power of the Atlas to provide information for genes known only by sequence. Furthermore, our analysis revealed several genes that were differentially expressed between these two clusters of putative hepatocytes, including *hmgcra*, *crp2* and *BX001030.1*, a poorly characterized gene (Figure 1B), demonstrating a molecular basis for cellular diversity among hepatocytes.

Next, we wondered whether these experiments captured rare cell types. Zebrafish embryos each have only about 30 primordial germ cells (PGCs) from 1 to 5 dpf (Tzung et al., 2015), thus providing a useful test case for detecting rare cells. We found that one computationally defined cluster (cluster 219) expressed the PGC markers *ddx4, nanos3, dnd1*, *and piwi2* (Houwing et al., 2008, 2007; Köprunner et al., 2001; Pandey et al., 2018; Weidinger et al., 2003; Yoon et al., 1997) indicating that PGCs were profiled and clustered in the Atlas (Figure 1C). The recovery of PGCs suggests that we have likely profiled most cell types present at the developmental stages analyzed in this study and provides an estimate of the lower threshold for detection of rare cell types. This finding does not rule out the possibility that some cell types may fail to be detected due to lack of dissociation, cell death, or inadequate sample size.

Finally, we sought to annotate all 220 clusters within the Atlas. We first identified the top 16 most differentially expressed genes for every cluster as having the highest ratio of expression among cells within a cluster relative to all other cells in the Atlas. Using RNA *in situ* expression patterns found in public databases, particularly ZFIN, we annotated the most likely cell type and corresponding tissue for each cluster (Fig 1A). These annotations occupy a nested hierarchy that contains information about germ layer, tissue type, and ultimately cell type (Table S2). Both unambiguous marker genes, with clear gene expression patterns referenced in ZFIN, as well as genes with no information on gene expression patterns appeared within the top 16 most differentially expressed genes for each cluster (Table S2). For most clusters (169/220), genes that have little or no previous expression information (genes beginning with “si:” or “zgc:” on ZFIN) appeared among the top sixteen most differentially expressed genes with an average of two (mean = 2.2, range = 0:9, SD = 2.1, for all clusters, (Table S2). For example, in the liver, notochord, and retina, the genes *si-dkey-96f10.1*, *si-ch1073-45j12.1*, and *si-dkey-16p21.8* were differentially expressed (respectively) and represent putative marker genes for these organs (Table S2). Thus, marker gene expression, coupled with analysis of RNA expression patterns *in vivo*, is an efficient means to annotate discrete clusters within the Atlas and provides a resource for helping to assign cell type information to poorly characterized genes.

### Combinatorial codes of gene expression inform anatomical position and identity

A critical aspect of any Atlas is that it should provide spatial information. This is a major challenge in scRNA-seq data derived from whole zebrafish because the anatomical position of any profiled cell is lost upon dissociation. For some cells, such as hepatocytes (see above), transcriptional profiles are distinct, and their anatomical location in the animal is restricted, making their identification relatively simple. However, many tissues, including the nervous system which serves as an extreme example due to its staggering functional and cellular diversity, present unique challenges because many of the cell types are transcriptionally similar, yet occupy anatomically disparate locations in the animal. For example, excitatory and inhibitory neurons reside throughout the nervous system, and the neurotransmitters and vesicular transporters that distinguish cell-types demonstrated broad expression across many discrete neural clusters in the Atlas (Figure S4), similar to what has been found in mouse (Cembrowski and Menon, 2018). We hypothesized that genes with spatially restricted expression domains in the anterior/posterior (A/P) and dorsal/ventral (D/V) axes would provide information about the subtypes of cells within the Atlas.

To examine this hypothesis, we used Atlas clusters that corresponded to the nervous system (Fig. 2A). We found these clusters had gene expression profiles that were associated with neural stem cells, differentiating, post-mitotic neurons, and mature neurons (Fig. 2B, Table S2) (Good, 1995; Ito et al., 2010; Wei et al., 2013) Within this subset of clusters from the Atlas, we examined genes known to have spatially restricted expression profiles along the A/P axis of the central nervous system and found that the clusters were organized in UMAP space in a manner that mimicked the *in vivo* A/P relationship of these genes (Fig. 2C) (Amores et al., 1998; Duggan et al., 2008; McClintock et al., 2001; Mori et al., 1994; Pandey et al., 2018; Scholpp et al., 2007; Scholpp and Brand, 2001; Woolfe and Elgar, 2007). The nervous system is well known for integrating both A/P and D/V information to generate particular cell fates. One of the best studied examples is spinal cord motorneuron specification where the so-called pMN ventral progenitor cells within the neural tube receive high levels of Sonic Hedgehog signaling turning on transcription factors such as *hoxa9a, olig2, isl1, isl2a, and mnx1* (Hutchinson and Eisen, 2005; Ravanelli and Appel, 2015). To identify spinal cord motorneurons we first used *hoxa9a* (Figure 2D) which is restricted to the posterior spinal cord (Prince et al., 1998) and thus distinguishes spinal cord neural cells. Next, we examined pMN markers that result from positional information from the D/V axis and found that they were expressed in increasingly restricted regions of the nervous system (Figure 2D). Strikingly, only a single cluster of neural cells mapped to the intersection of the A/P and D/V markers (Fig. 2A,D – magenta arrow head, and 2E), and this cluster expresses genes such as *slc18a3a*, the acetycholine neurotransmitter transporter, *mnx1*, a motor neuron marker gene, and *snap25a*, which marks differentiated neurons (Bayés et al., 2017; Nakano et al., 2005; Wei et al., 2013). These results indicate that a combinatorial code of spatially confined transcription factors with motoneuron fate determining molecules is sufficient to identify anatomically and functionally distinct populations of neurons in the Atlas. This approach was applied to all central nervous system clusters of the Atlas (Table S2) and provides a roadmap to uncover the regional transcriptional differences amongst anatomically and functionally diverse neurons. We conclude that spatially restricted gene expression can be mapped to the Atlas even given complicated and similar transcriptional profiles, enhancing our knowledge of cell-type-specific gene expression within these domains.

**Figure 2.**
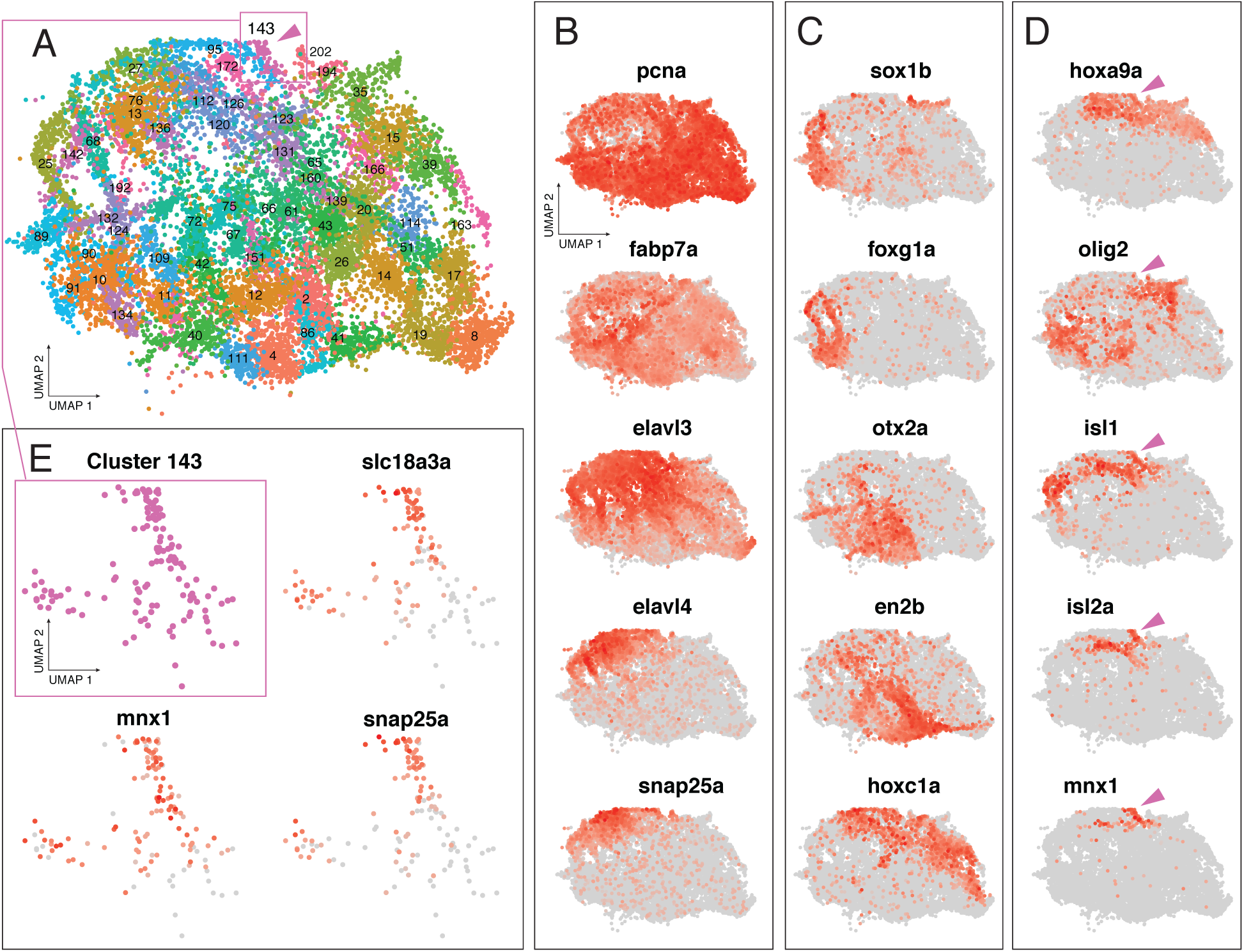
Spatially restricted gene expression patterns mapped to the Atlas. (A) Clusters from the Atlas with annotated neural cell identifies. The cells represented correspond to the dashed oval in Figure 1A. Cluster 143 is marked by magenta arrowhead. Magenta box describes clusters enlarged in E. (B) Neural maker genes show regions of progenitor (*pcna*, *fabp7a*) and differentiated neural identities (*elavl3*, *elavl4*, and *snap25a*). (C) Spatially restricted transcription factors show discrete domains of expression across neural clusters in the Atlas. Cells with expression of genes related to the forebrain/telencephalon (*sox1b*, *foxg1a*), midbrain and hindbrain (*otx2a*, *en2b*), and anterior spinal cord (*hoxc1a*), cluster together in UMAP space. (D) Combinatorial code of gene expression within *hoxa9a* expressing domain reveals cluster of putative spinal cord motor neurons expressing markers of the progenitor domain (*olig2*) and differentiation (*isl1, isl2a, mnx1*). Cluster 143 marked by magenta arrowhead. (E) Magnified view of cluster 143 shows high levels of motor neuron marker genes *slc18a3a* and *mnx1* and the differentiated neuronal marker *snap25a*.

### Cell-type specific gene expression changes over time are mapped in the Atlas

A primary goal of this Atlas is to provide temporal information about gene expression changes associated with zebrafish development across four days of development. This approach is critical for understanding the genetic basis of how stem cells and progenitor cells produce diverse progeny, and the pathways along which differentiated cells mature during organogenesis to form functioning tissues.

By aggregating our replicate experiments from three developmental stages, we sought to determine which changes in gene expression were associated with cell-type specification and organogenesis during this period of development. Within the transcriptional mapping space of the Atlas, cell clusters often separated according to developmental age (Fig. 3A). Although a major discretizing factor across the Atlas is time of origin, within any particular transcriptionally identified cell type there are examples of apparent transcriptional continuity that connect developmental timepoints. For example, the notochord formed several discrete clusters across developmental time (Figure 3B,C). We identified notochord cells based on the exclusive expression of *tbxta* (Odenthal et al., 1996), and in addition, high levels of *col2a1a, col9a2 and matn4* expression help verify notochord cluster identity (Figure S5) (Garcia et al., 2017; Halpern et al., 1997; KO et al., 2005), suggesting that cells of the notochord had been accurately clustered together. Thus, the Atlas has effectively clustered cells of the notochord with one another in UMAP space and has also preserved transcriptional differences associated with developmental time.

**Figure 3.**
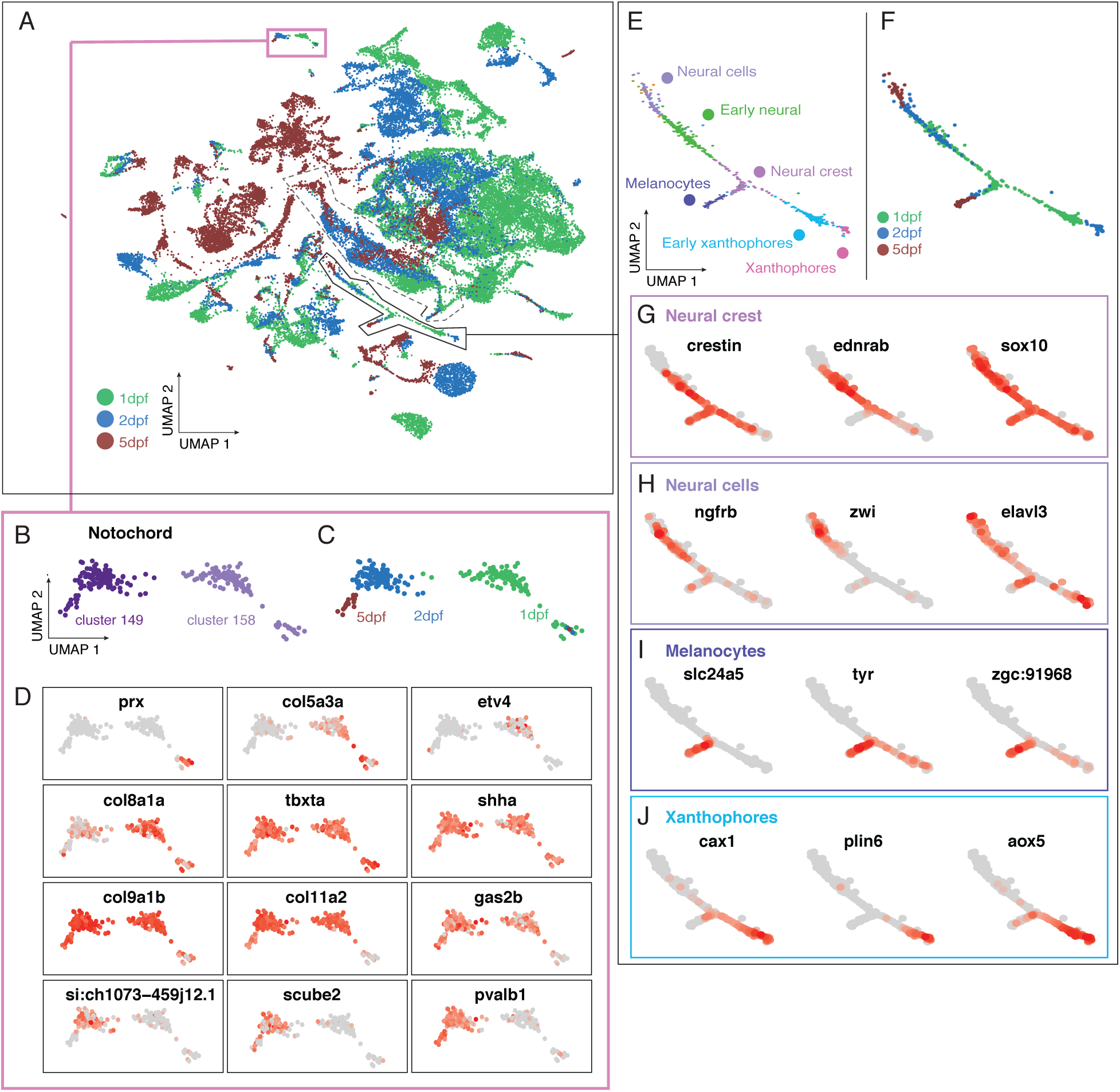
Temporal gene expression analysis in the Atlas using cell age. (A) Aggregating cells from 1, 2, 5 dpf embryos reveals age-related changes in gene expression for discrete clusters. Pink box describes notochord clusters enlarged in B-D; solid black line shows approximate region of neural crest lineages enlarged in E-J. Dashed line shows approximate region of retinal cells described in Figure 4. (B-C) Notochord clusters contain cells of distinct ages. (D) Differential gene expression analysis in developing the notochord. (E) Discrete clusters corresponding to cell type specification in neural crest lineages are annotated with cell-type and color coded by cluster. (F) Age of cells associated with neural crest lineage. (G-J) Differential gene expression analysis over time in neural crest progenitors (G), neural cells (H), melanocytes (I), and xanthophores (J) across developmental time.

Next, we asked which genes were differentially expressed between notochord clusters of progressing developmental age. We found *col5a3a, prx*, and *etv4* to have enriched expression in 1 dpf notochord, a poorly characterized gene*, si-ch1073-45j12.1* expressed in 2 dpf notochord, but repressed at 5 dpf, while *scube2, pvalvb1, gas2b* were specifically expressed in older, 2-5 dpf notochord (Figure 3D). Importantly, we identified genes that are expressed in notochord throughout the developmental stages analyzed in this study (common to both notochord clusters), including *col9a1b, col11a2*, and *shha* (Figure 3D). Mining these data suggests functional experiments to understand the mechanisms that drive notochord differentiation. We conclude that temporal changes in gene expression are efficiently captured in the Atlas by comparing transcriptional profiles from cells of the same tissue at progressing developmental ages.

Next, we reasoned that the Atlas would provide access to unique transcriptional trajectories of cell type specification in developmental time in related cell types. A striking example appeared in the neural crest lineage (Fig. 3E,F). Clusters corresponding to neural crest progenitors were identified using *crestin, ednrab*, and *sox10* (Figure 3G) (Bonano et al., 2008; Dutton et al., 2001; Luo et al., 2001). Neural progeny of the neural crest were characterized by expression of *ngfrb, zwi*, and *elavl3* (Schaefer and Brösamle, 2009), melanocytes by *slc24a5* and *tyr* expression (Braasch et al., 2009, 2007; Kelsh et al., 2000), and xanthophores by *cax1, plin6*, and *aox5* expression (Figure 3G-J) (Granneman et al., 2017; Manohar et al., 2010). We were immediately struck by the distinct transcriptional profile corresponding to these differentiated cell-types as well as their apparent similarity, due to their proximity in UMAP coordinates, to the neural crest precursors that produce them (Figure 3E). The continuous nature of the transcriptional state of this lineage led us to ask whether the developmental stage of each cell type (age of embryo used in this study) corresponded to its position within the UMAP-derived lineage. Marking the developmental stage of embryos used in each experiment unambiguously resolved that cells derived from older embryos were associated with the most differentiated state of each apparent developmental trajectories (Figure 3F). Furthermore, the neural crest precursors that produce these cells were primarily obtained from younger embryos (Figure 3F). Interestingly, a poorly described gene, *zgc:91968*, demonstrated high levels of expression in the melanocyte cluster (Figure 3I and Table S2) confirming recent expression analysis of this gene (Heffer et al., 2017), and *si:dkey-25li10*.2 and *si:dkey-129g1*.9 in xanthophore clusters (Table S2), expanding our knowledge of cell-type specific expression patterns for these transcripts. Thus, the Atlas is a useful tool for investigating gene expression changes associated with the differentiation of multiple cell types from a lineage of common progenitors.

### Pseudotemporal analysis using Monocle bolsters temporal gene expression analysis during development

To validate changes in gene expression identified by comparing cells of similar types but from different developmental stages, we focused on developing retinal lineages as they provide striking example (Figure 4A,B). We identified clusters of retinal pigment epithelium based on expression of *cx43* (Figure 4C) (Dermietzel et al., 2000); retinal progenitors using *vsx2 and col15a1b* (Figure 4D) (Pujic et al., 2006; Vitorino et al., 2009); differentiating retinal cells using *crx* and *neurod1* (Hitchcock and Kakuk-Atkins, 2004; Ochocinska and Hitchcock, 2009, 2007; Shen and Raymond, 2004) and also found differential expression of *otx2b, foxn4, hes2.2 and ntoh7* (Figure 4E); retinal neurons using *lin7a* (Wei et al., 2006) and also found differential expression of *alcama* and *barhl2* (Figure 4F); photoreceptors using *rho* (Vihtelic et al., 1999) and also found differential expression of *grk1a and opn1lw2* (Figure 4G). Interestingly, the proximity in UMAP coordinates of these cell types suggested that their transcriptomes were related in a continuous fashion (Figure 4A), similar to the notochord and neural crest lineages (see above). Furthermore, the youngest cells (1 dpf) were strongly enriched within the retinal progenitor cluster, whereas photoreceptors and retinal neuron clusters mostly contained older cells (5 dpf) (Figure 4A-B). Between these progenitors and differentiated cells, we found cells primarily from 2 dpf embryos (Figure 4B). While there was a general progression from younger to older cells across the UMAP space, we also found cells derived from 2 and 5 dpf larvae within the clusters defined as retinal progenitors, and 5 dpf cells within the retinal differentiation clusters (Fig. 4A-B). This was expected as retinal development continues throughout life (Malicki et al., 2016) and there is a large amount of proliferation and specification occurring over the time period assayed in our experiments.

**Figure 4.**
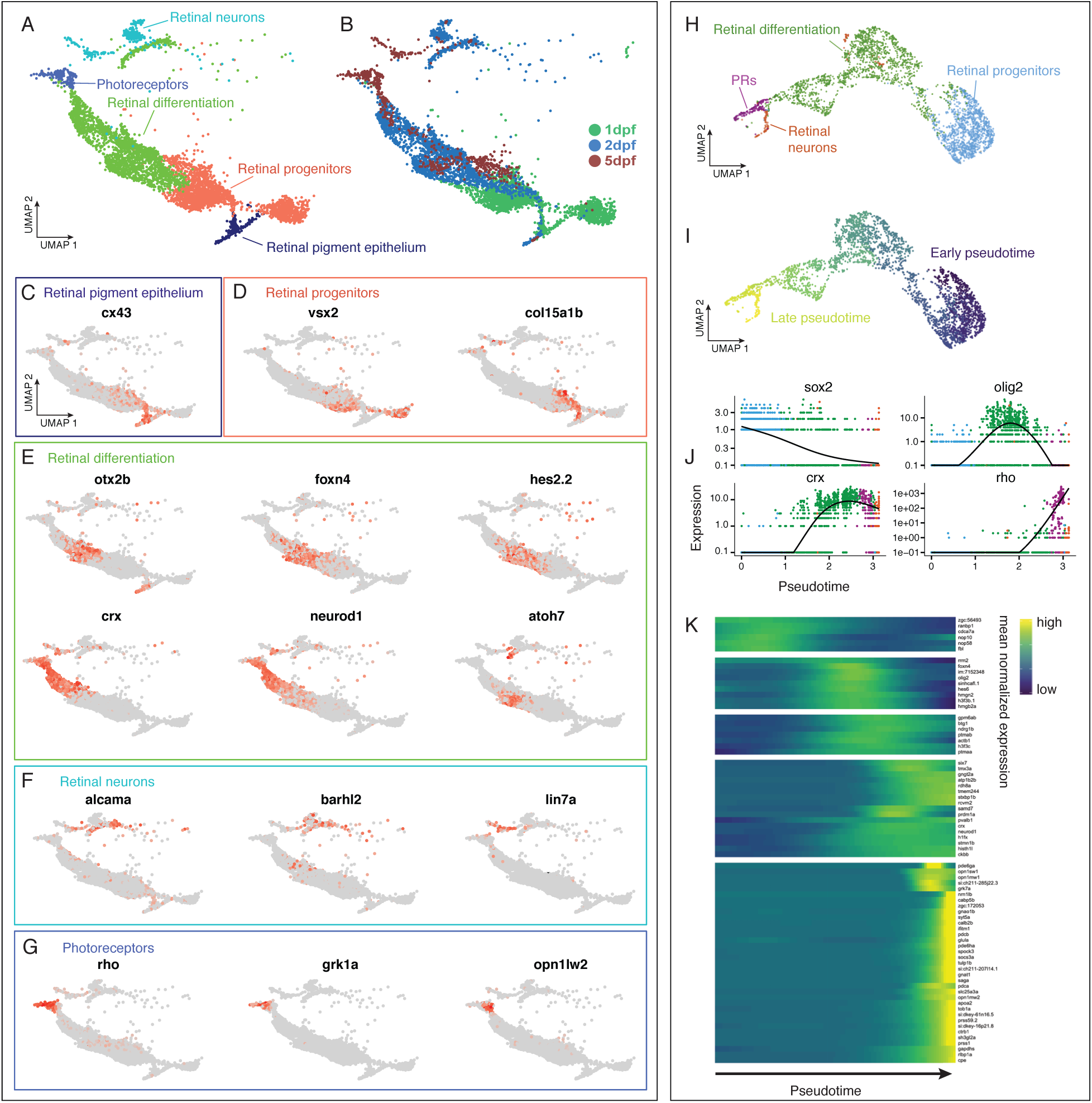
Temporal and pseudotemporal gene expression analysis of the retina. (A) Retinal cells form continuous transitions in UMAP space. Colors correspond to groupings of clusters and annotations with the cells represented corresponding to the dashed oval in Figure 3A. (B) Age of cells associated with retinal cell clusters. (C-H) Differential expression analysis over time during retina development in clusters of putative (C) retinal pigment epithelium, (D) retinal progenitors, (E) differentiating retinal cells, (F) retinal neurons, and (G) photoreceptors. (H-K) Pseudotemporal analysis of retina neuron development in Monocle. (H) Monocle reconstruction of retinal neural lineages agrees with UMAP analysis and annotation (compare to A). (I) Monocle reconstruction of retinal lineage colored by pseudotemporal age. (J) Monocle analysis reveals progressive repression of *sox2*, activation of *crx* and *rho*, and transient activation of *olig2* across pseudotime. (K) Heatmap of gene expression analysis associated with retinal cell type specification in Monocle.

We sought to compare relationships based on known ages to an unsupervised, pseudotemporal analysis of changes in gene expression across the retinal lineage. We omitted the clusters associated with retinal pigment epithelium from Monocle analysis, as these cells differentiate from retinal progenitors independently of retinal neurons (Fuhrmann et al., 2014). This analysis using Monocle (Cao et al., 2019) revealed many *de novo* predicted genes associated with retinal development, and confirmed lineage relationships among cells from distinct developmental stages. Strikingly, the unsupervised pseudotemporal analysis strongly correlated with the developmental stage of each cell type within the retinal lineages (Figure 4H). Specifically, Monocle described a pseudotemporal order of retinal cells with striking fidelity to our annotations, and in concert with their developmental age (compare Figure 4H with Figure 4A-B). This result suggests that temporal changes in gene expression occur in a continuous fashion from progenitor to differentiated photoreceptors and retinal neurons. This pseudotemporal analysis helped identify genes that are downregulated during retinal differentiation, such as *sox2* and also genes that are upregulated during retinal differentiation, such as *crx, olig2 and rho* (Figure 4J). Additional suites of genes with temporally restricted expression occurring throughout retinal differentiation were identified by Monocle providing a rich resource for understanding genes associated with retinal progenitors and photoreceptor specification during development (Figure 4K). These results highlight the usefulness of the Atlas for investigating temporal changes in gene expression during zebrafish development.

## DISCUSSION

The Atlas presented here is a tool for researchers to investigate gene expression, developmental trajectories, and identify cell types in developing zebrafish. Our profiling of 44,102 single cell spanning one to five days post fertilization provide transcriptional insight into how all of the major organs form in this vertebrate system. We annotated all 220 bioinformatically identified clusters and highlighted several strategies for interrogating changes in gene expression associated with the development of zebrafish embryos at single cell resolution. Specifically, we demonstrate that: 1) algorithmically derived clusters can be annotated using marker gene expression, 2) expression analysis can advance knowledge for genes previously known only by sequence, and 3) temporal gene expression changes track continuous changes in gene expression patterns during differentiation of specific cell-types.

Recent work has investigated single-cell transcriptional changes during earlier stages in zebrafish development, specifically embryogenesis (up to 24 hpf) (Farrell et al., 2018; Wagner et al., 2018). These data, and the ever-increasing body of cell-type and transcriptomic data, represent some of the first system-level molecular and genetic insight to be leveraged toward understanding animal development. Technical advances that enable broader coverage of the single-cell transcriptome, reduce the cost associated with these experiments, and computational annotation algorithms to annotate cell type based on independent expression databases will accelerate their contribution to developmental biology and medicine. We, and other researchers, are increasingly aware that a new age of developmental biology is emerging in which we must pair a reductionist approach of analysis of specific mechanisms of function with an understanding of system-level effects. We view a comprehensive Atlas as a first step towards such an understanding, and believe the knowledge gained from experimentation in easily modifiable model organisms such as zebrafish will greatly enhance efforts toward unifying and translational applications of biology.

The clusters presented in this study require additional work to improve the accuracy and utility of this Atlas. Specifically, researchers with deep expertise regarding cell-type specific gene expression can help modify our tentative cluster annotations, which will improve this initial version, thus making it more useful for hypothesis testing. We envision this as an ongoing process with the community and make the current raw data, analysis methods, and our current draft of cluster identification available to the community (https://biology.uoregon.edu/profile/acmiller), with ongoing work and feedback continually improving the utility of the Atlas going forward. Furthermore, we view gene expression analysis in the Atlas as a hypothesis-generating process that will require subsequent functional experiments to test the accuracy of the *in silico* predictions presented here. For example, our Atlas has revealed novel cell and developmental stage-specific gene expression profiles that require RNA *in situ* hybridization to validate expression in a predicted cell type, and also loss of function analysis to examine their function in development. The results of such experiments will be forthcoming from our group and we anticipate results from other research groups, as well. Furthermore, our ongoing goal is to develop a Zebrafish Cell Atlas, spanning all stages of zebrafish development at fine temporal and cellular resolution that will be made freely available through ZFIN. We envision that this will provide a framework to answer questions in all areas of vertebrate developmental biology and provide an open resource for the community to accelerate discovery.

## METHODS

Fish were maintained by the University of Oregon Zebrafish facility using standard husbandry techniques (Westerfield, 2007). Embryos were collected from natural matings, staged and pooled (n=15 per replicate). Animals used in this study were: *Tg(olig2:GFP)vu12* for 1 dpf, n=2; 2 dpf, n=1; 5 dpf, n=2 samples and *Tg(elavl3:GCaMP6s*) for 2 dpf, n=1 sample (Figure S1).

### Embryo dissociation

Collagenase P was prepared to a 100mg/mL stock solution in HBSS. Chemical dissociation was performed using 0.25% Trypsin, Collagenase P (2mg/mL), 1 mM EDTA (pH 8.0), and PBS for 15min at 28C with gently pipetting every 5 min. Dissociation was quenched using 5% calf serum, 1mM CaCl2, and PBS. Cells were washed and resuspended in chilled (4C), 1% calf serum, 0.8 mM CaCl2, 50 U/mL penicillin, 0.05 mg/mL streptomycin, and DMEM and passed through a 40uM cell strainer (Falcon) and diluted into PBS + 0.04% BSA to reduce clumping. A final sample cell concentration of 2000 cells per microliter, as determined on a Biorad TC20 cell counter, was prepared in PBS + 0.04% BSA for cDNA library preparation.

### Single-cell cDNA library preparation

Sample preparation was performed by the University of Oregon Genomics and Cell Characterization core facility (https://gc3f.uoregon.edu/). Dissociated cells were run on a 10X Chromium platform using 10x v.2 chemistry targeting 10,000 cells. Dissociated samples for each time point (1, 2 and 5 dpf) were submitted in duplicate to determine technical reproducibility. Replicates for 1 and 5 dpf samples were from different clutches, but prepared in tandem, on the same day. Replicates for 2 dpf samples were prepared from different transgenic backgrounds on different days. The resulting cDNA libraries were amplified with 15 cycles of PCR and sequenced on either an Illumina Hi-seq (5/6 samples) or an Illumina Next-seq (n=1, 48h dpf sample) (Figure S1).

### Computational analysis

The resulting sequencing data were analyzed using the 10X Cellranger pipeline, version 2.2.0 (Zheng et al., 2017) and the Seurat (Satija et al., 2015) software package for R, v3.4.4 (R Development Core Team, 2011) using standard quality control, normalization, and analysis steps. Briefly, we aligned reads to the zebrafish genome, GRCz11_93, and counted expression of protein coding reads. The resulting matrices were read into Seurat where we performed PCA using 178 PCs based on a Jack Straw determined significance of P < 0.01. UMAP analysis was performed on the resulting dataset with 178 dimensions and a resolution of 13.0, which produced 220 clusters and one singleton. Differential gene expression analysis was performed using the FindAllMarkers function in Seurat. Cells were subsetted based on gene expression and cluster for Monocle analysis (see below). We compared the correspondence between our replicates by counting the number of cells from each sample, in every cluster, and plotting these values in a scatterplot (x,y = fraction of cells from sample in a single cluster from replicate one, fraction of cells in a single cluster from replicate two) and performed a regression analysis.

### Pseudotemporal Trajectory analysis

We subsetted the retinal progenitors, retinal neurons and photoreceptor cells and applied UMAP dimensionality reduction to project cells in three dimensions using the preprocessCDS and reduceDimension functions in Monocle (v.2.99.1) (Cao et al., 2019) using default parameters (except for preprocessCDS: num_dim = 20; reduceDimension: reduction_method=UMAP, metric=cosine, n_neighbors=20, mid_dist=0.2). To learn the pseudotemporal trajectory, we then used the partitionCells, learnGraph, and orderCells functions using default parameters (except for partitionCells: k = 15; learnGraph: close_loop = TRUE, ncenter=400). To determine differentially expressed genes over pseudotime, we filtered the data set for genes expressed in at least 5 cells and performed differential expression analysis using a full model of sm.ns(Pseudotime, df=3). The top 80 DEGs were selected by q-value and plotted using the plot_pseudotime_heatmap function in Monocle.

### Cluster annotation

We identified the top 16 most differentially expressed genes for every cluster as having the highest ratio of expression among cells within a cluster relative to all other cells in the Atlas. Using RNA *in situ* expression patterns found in public databases, particularly ZFIN, we annotated the most likely cell type and corresponding tissue for each cluster. These annotations occupy a nested hierarchy that contains information about germ layer, tissue type, and ultimately cell type (Table S2).

## Author contributions

DF performed all experiments, analyzed data and co-wrote the manuscript. LS performed pseudotime analysis in Monocle. AM analyzed data and co-wrote the manuscript.

## Acknowledgements

We thank Charles Kimmel, Monte Westerfield, Kryn Stankunas, Judith Eisen and John Postlethwait for many helpful conversations during the conceptualization of the project and preparation of the manuscript. We thank the University of Oregon Fish Facility for superb animal care. Funding was provided by the NIH Resource-Related Research Project Grant R24OD026591 and the University of Oregon to A.C.M.

---

**Figure S1.** Summary of statistics related to 10x Chromium cDNA library preparation and sequencing. (A) The sample name, genotype, age, reads per cell and sequencing platform used for individual experiments. (B) Summary of read, transcript, gene and cellular diversity and abundance for aggregated data used to produce the Atlas.

Replicate experiments for 1 dpf (C), 2 dpf (D) and 5 dpf (E) show mutual representation across clusters and indicate high reproducibility. We analyzed the correspondence between our replicates by counting the proportion of cells in each independent replicate, in every cluster and plotting these values in a scatterplot (x,y = proportion of all cells in a single cluster from replicate one, proportion of all cells in a single cluster from replicate two) and found a high level of correlation with R^2^ values of 0.8 (1 dpf), 0.4 (2 dpf) and 0.8 (5 dpf).

**Figure S2.** UMAP plot of all 220 clusters in the Atlas. Replicate experiments from 1, 2 and 5 dpf embryos are aggregated into a single plot. Cluster numbers are applied to centroid of each cluster and correspond to Table S1 and Table S2 and cluster names referenced throughout this work.

**Figure S3.** Heatmap representation of genes represented in the first principal component (PC1) following PCA analysis. The hepatocyte markers *cp* and *fabp10a* are among the most differentially expressed genes. Yellow = high expression, magenta = low expression. 500 cells are plotted.

**Figure S4.** Genes encoding neurotransmitter vesicular transporters and biosynthesis pathways are markers of neuronal subtypes including, *slc17a6a / slc17a6b* for glutamatergic neurons, and *gad1a / gad1b* for gamma-aminobutyric acid (GABA), GABAergic neurons. Red = high expression, gray = low expression.

**Figure S5.** Marker genes *tbxta*, *matn4*, col9a2, and *col2a1a*, enable annotation of discrete clusters of notochord cells. Red = high expression, gray = low expression.

**Table S1.** List of differentially expressed genes for each cluster. The table was generated using the FindAllMarkers command of Seurat.

**Table S2.** The top 16 most differentially expressed genes for each cluster that were used for annotation. Our annotation includes the germ layer, tissue and cell type that correspond to each cluster. Also noted is the presence of cells from each of the developmental time points profiled in this study (1,2 and 5 days post fertilization). These parameters are displayed as column headings.

