## Supplemental Figures and Tables for "A Single-Cell Transcriptome Atlas for Zebrafish Development"

### Figure S1

A

| Sample | Genotype | Age | Reads per cell | Sequencer |
|---|---|---|---|---|
| 1a | *Tg(olig2:GFP)vu12* | 1 dpf | 62,016 | HiSeq 4000 |
| 1b | *Tg(olig2:GFP)vu12* | 1 dpf | 49,699 | HiSeq 4000 |
| 2a | *Tg(elavl3:GCaMP6s)* | 2 dpf | 74,928 | NextSeq 500 |
| 2b | *Tg(olig2:GFP)vu12* | 2 dpf | 53,287 | HiSeq 4000 |
| 5a | *Tg(olig2:GFP)vu12* | 5 dpf | 135,685 | HiSeq 4000 |
| 5b | *Tg(olig2:GFP)vu12* | 5 dpf | 122,920 | HiSeq 4000 |

B

| **Post-aggregation** | |
|---|---|
| Number of Cells | 44,102 |
| Fraction reads in cells | 71.5% |
| Mean reads per cell | 74,914 |
| Median genes per cell | 1,807 |
| Median UMI counts per cell | 8,345 |
| Total number of reads | 3,303,866,971 |

C - 1 dpf

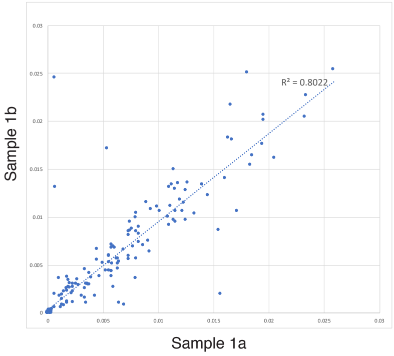


D - 2 dpf

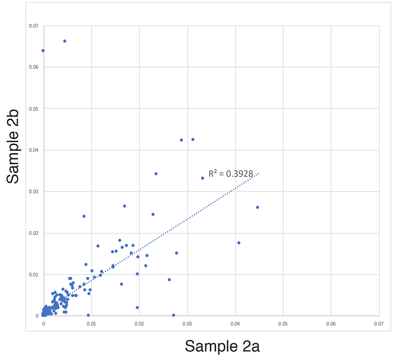


E - 5 dpf

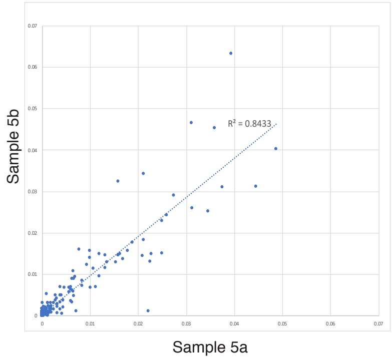

### Figure S2

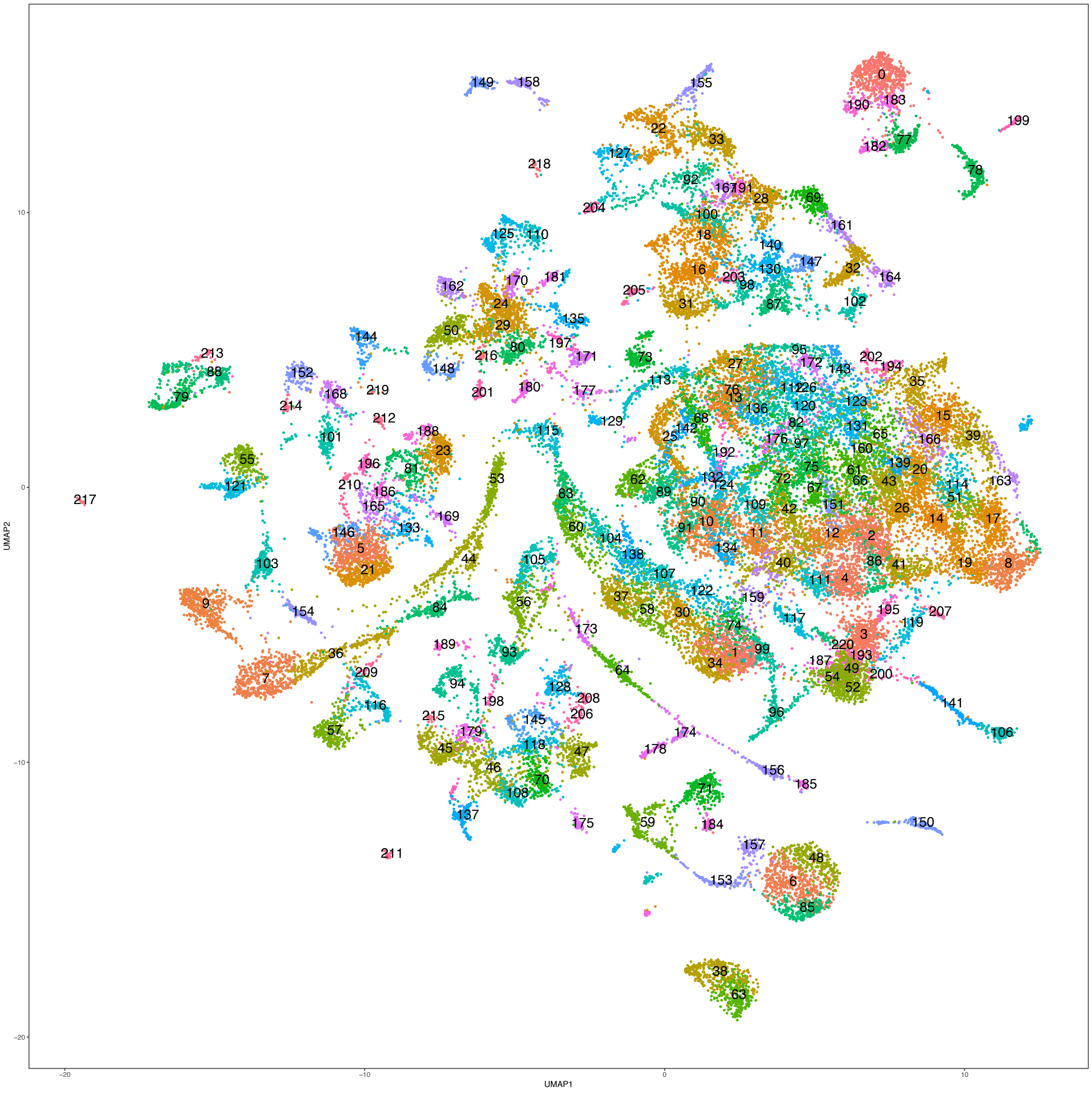

Figure S3

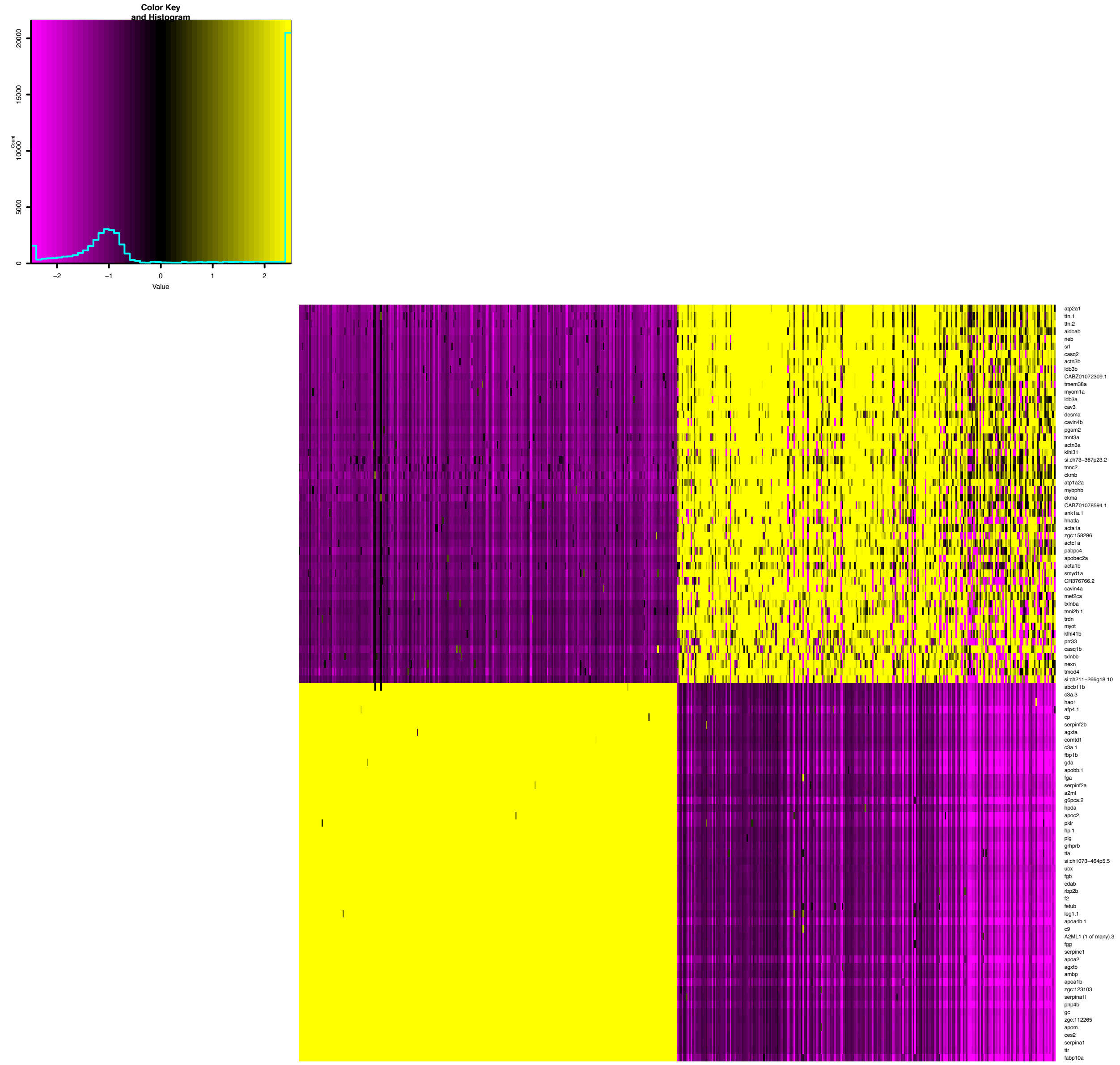
Color Key
and Histogram
Count
20000
15000
10000
5000
0
−2
−1
0
1
2
Value
atp2a1
ttn.1
ttn.2
aldoab
neb
srl
casq2
actn3b
ldb3b
CABZ01072309.1
tmem38a
myom1a
ldb3a
cav3
desma
cavin4b
pgam2
tnnt3a
actn3a
klhl31
si:ch73−367p23.2
tnnc2
ckmb
atp1a2a
mybphb
ckma
CABZ01078594.1
ank1a.1
hhatla
acta1a
zgc:158296
actc1a
pabpc4
apobec2a
acta1b
smyd1a
CR376766.2
cavin4a
mef2ca
txlnba
tnni2b.1
trdn
myot
klhl41b
prr33
casq1b
txlnbb
nexn
tmod4
si:ch211−266g18.10
abcb11b
c3a.3
hao1
afp4.1
cp
serpinf2b
agxta
comtd1
c3a.1
fbp1b
gda
apobb.1
fga
serpinf2a
a2ml
g6pca.2
hpda
apoc2
pklr
hp.1
plg
grhprb
tfa
si:ch1073−464p5.5
uox
fgb
cdab
rbp2b
f2
fetub
leg1.1
apoa4b.1
c9
A2ML1 (1 of many).3
fgg
serpinc1
apoa2
agxtb
ambp
apoa1b
zgc:123103
serpina1l
pnp4b
gc
zgc:112265
apom
ces2
serpina1
ttr
fabp10a

Figure S4

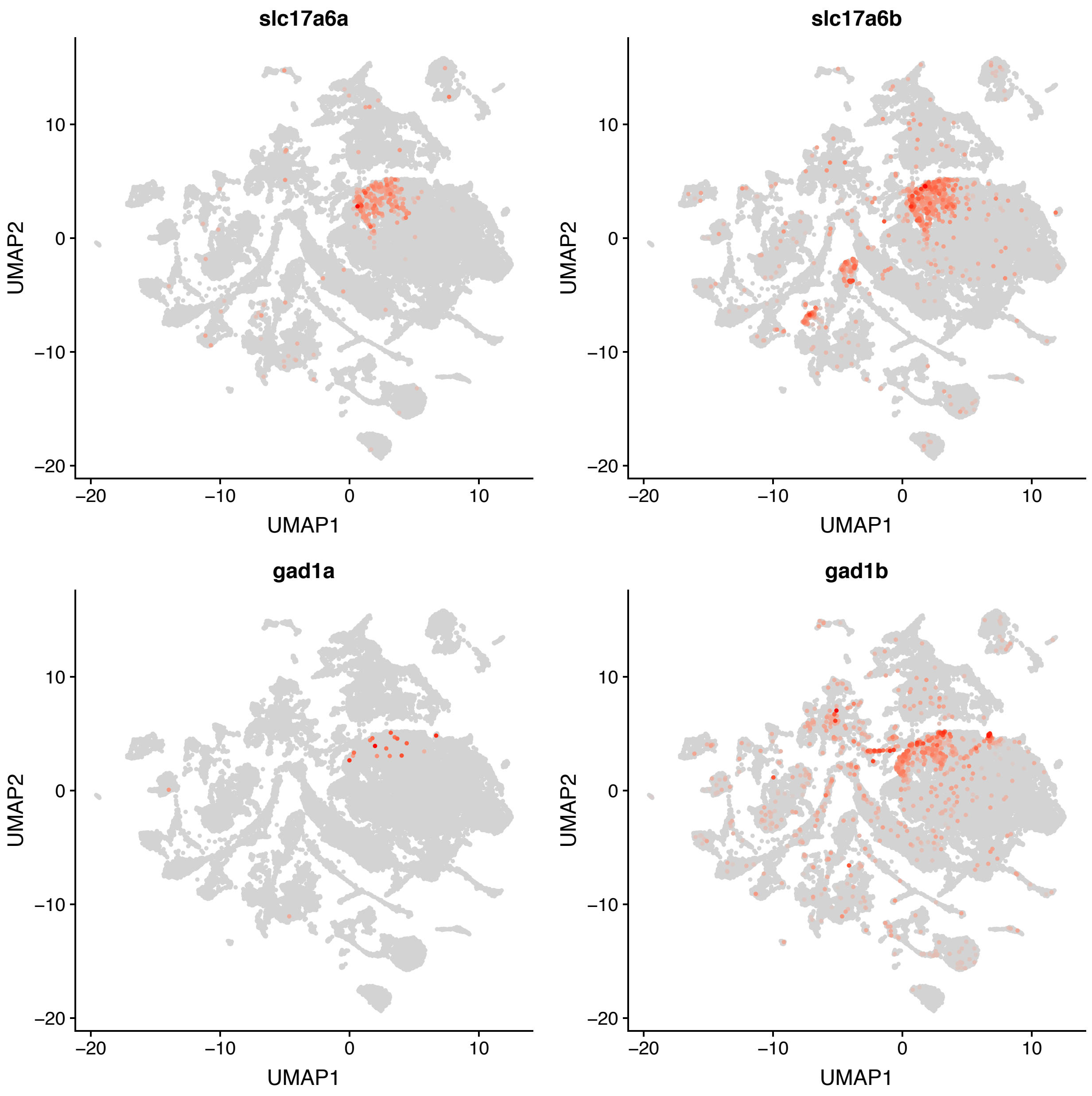

Figure S5

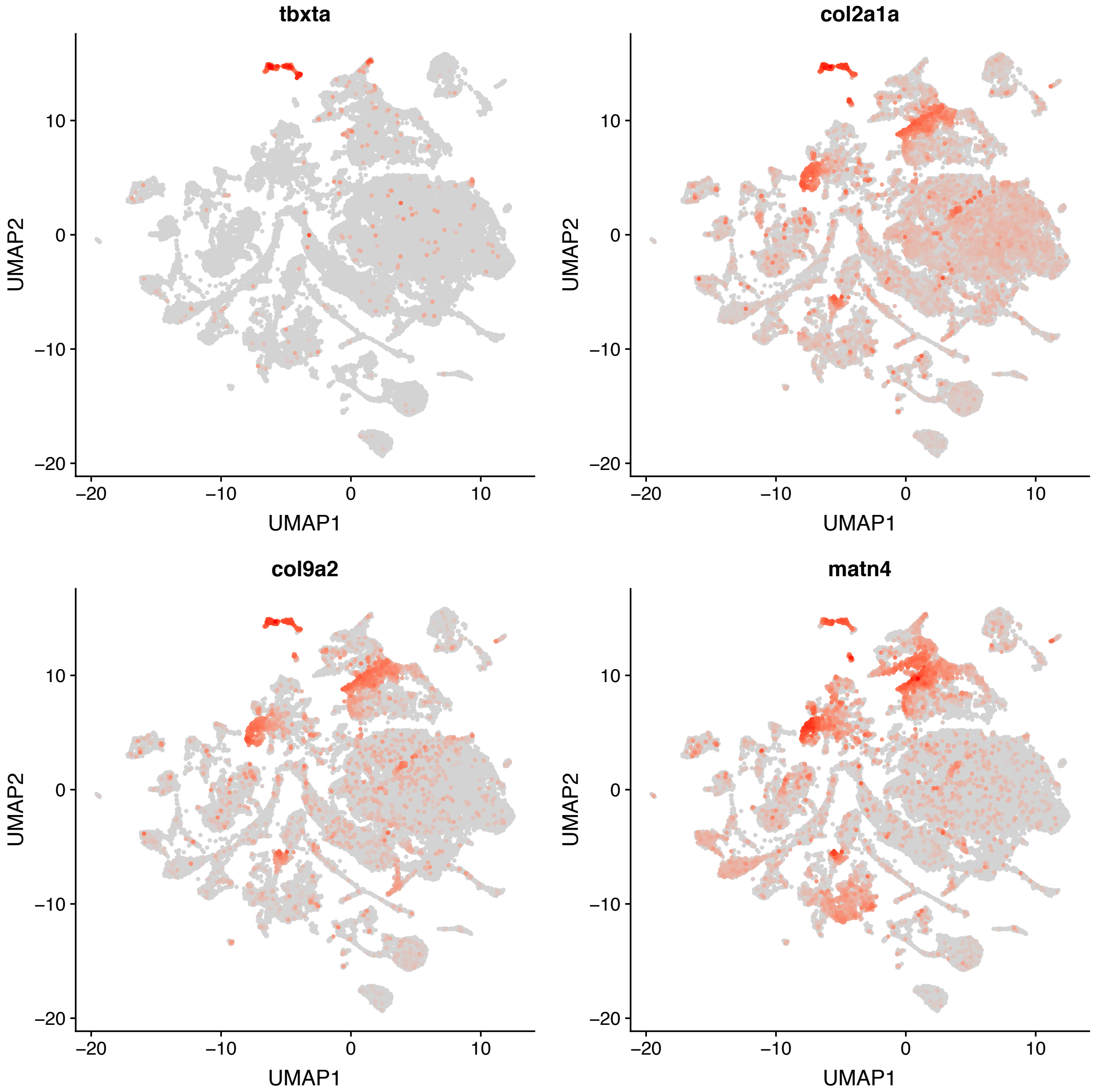

Table S1

https://biology.uoregon.edu/profile/acmiller/

### Table S2

| Cluster ID | Germ Layer | Tissue | Cell type | Subtype | Ident Annotation | Groupname | Day of Origin | Top16 most differential genes | | | | | | | | | | | | | | | | n=si+zgc genes |
|---|---|---|---|---|---|---|---|---|---|---|---|---|---|---|---|---|---|---|---|---|---|---|---|---|
| 0 | Ectoderm | Integument | Periderm | | Peri1a | Peri1 | 1 | he1.3 | ftr14l | cyp2aa8 | cyp2aa11 | si:ch211-264 | si:dkeyp-87d | si:dkey-160o | BX569789.1 | zgc:113442 | zgc:194839 | nos1 | tgm1l4 | CABZ010493 | si:ch211-202 | nadka | si:dkeyp-4c4. | 7 |
| 1 | Ectoderm | Central Nervous System | Neuroblast | Retinal | RetProgAlla | RetProg | 1,2,5 | vsx2 | rx1 | six3b | si:dkey-238o | col15a1b | stra6 | nr2e1 | myb | wfdc1 | tbx2a | crip3 | hmx4 | pax6b | cart3 | AL954322.2 | zic2b | 1 |
| 2 | Ectoderm | Central Nervous System | Neuron, differentiating | Mid-hindbrain boundary | MHBDiff12a | MHB | 1,2 | her3 | en2b | sfrp1a | sp8b | dbx1b | efna2a | dbx1a | wfikkn1 | sox21a | col5a3a | sema3fa | npas1 | st8sia2 | sfrp5 | irx1a | prtgb | 0 |
| 3 | Nonspecific genes | | | | NonspecificAlla | Nonspecific | 1,2,5 | en2b | sox19a | paics | cmss1 | her8a | esco2 | sox3 | nusap1 | umps | ftsj3 | rrm2 | pclaf | mcmbp | nasp | parp1 | ddx24 | 0 |
| 4 | Ectoderm | Central Nervous System | Neuroblast | Mid-hindbrain boundary | MHBProg12 | MHB | 1,2 | dmbx1a | pax7a | en2b | pax7b | en2a | efna2a | hey2 | otx2b | pax3a | sema3fa | foxn4 | otx2a | fzd10 | irx1a | mab21l2 | pou3f3a | 0 |
| 5 | Ectoderm | Integument | Integument Gill? | Gill | Gill5a | Gill | 5 | urgcp | plekhs1 | si:dkey-201i2 | AL928650.1 | si:ch211-266 | si:dkey-187j1 | ca15c | si:dkey-239b | si:dkey-238k1 | cyldb | CABZ010516 | vwa10.1 | CU914622.2 | rhcgl1 | zgc:194007 | si:ch73-329n | 7 |
| 6 | Mesoderm | Blood | Hematopoietic cell | Nucleate erythrocyte | RBC2a | RBC2 | 2 | gp9 | rfesd | hif1al2 | zgc:163057 | si:ch211-207 | si:ch211-250 | si:dkey-93h2 | si:ch73-250a | nmt1b | hbae5 | si:ch211-227 | epb41b | alas2 | rell2 | CR293511.1 | nt5c2l1 | 6 |
| 7 | Mesoderm | Muscle | Skeletal muscle | Fast muscle cell | MusSkelFast1 | MusSkelFast | 1 | rtn2a | obsl1a | klhl40b | prp | dhrs7ca | p2rx5 | klhl43 | neurl2 | FO818704.2 | zgc:113276 | xirp2b | obscnb | tgm2a | slc8a3 | CU929145.2 | cacna1sb | 1 |
| 8 | Ectoderm | Central Nervous System | Neuroblast | Hindbrain | HBProg1a | HB | 1 | atoh1a | olig4 | olig3 | egr2b | nog2 | cdh6 | msx3 | epha4l | fibinb | atoh8 | msx1b | zic2b | irx1b | bmp5 | draxin | rfx4 | 0 |
| 9 | Mesoderm | Muscle | Skeletal muscle | Fast muscle cell | MusSkelFast2a | MusSkelFast | 2 | hspb2 | capn3b | CABZ010615 | pbld2 | hsc70 | pfkma | tmem41ab | glra4a | igfn1.3 | synpo2b | pfkmb | CU693479.1 | kbtbd12 | cdnf | myha | abrab | 0 |
| 10 | Ectoderm | Central Nervous System | Neuroblast | Forebrain | FBProgAlla | FB | 1,2,5 | foxg1a | fezf1 | gsx2 | emx3 | fezf2 | sox1b | lhx2b | lmo1 | cyp26c1 | CR848047.1 | LO018196.1 | mcama | six3a | auts2b | CU467822.1 | zgc:165461 | 1 |
| 11 | Ectoderm | Central Nervous System | Neuron, differentiating | Midbrain | MBDiff5 | MB | 5 | en2a | fsta | atp1b4 | slc3a2a | znf219 | atp1a1b | BX324164.4 | ctrl | c6ast4 | opn1sw1 | cpa5 | ela2l | gngt2b | ptgdsb.2 | arr3b | ctrb1 | 0 |
| 12 | Ectoderm | Central Nervous System | Neuroblast | Midbrain | MBProg12a | MB | 1,2 | shox2 | otx2a | zgc:101100 | dbx1a | barhl2 | CR751602.2 | robo4 | nr2e1 | foxb1a | otx2b | tcf7l2 | pou3f2b | irx1b | BX569787.1 | otx1 | mapk12b | 1 |
| 13 | Ectoderm | Central Nervous System | Neuron | Mid-hindbrain boundary | MHBNeur25 | MHB | 2,5 | dpf1 | pou4f1 | kif3cb | diras1a | sprn | ppp1r14ba | si:ch211-10a | atl1 | camk2d2 | si:dkey-12h9. | nell2b | nmnat2 | si:dkey-56f14 | BX927184.1 | si:dkeyp-117 | atat1 | 4 |
| 14 | Ectoderm | Central Nervous System | Neuroblast | Hindbrain | HBProg1b | HB | 1 | lbx1b | gsx1 | pax7b | si:ch73-364h | pax7a | sox21a | prdm8 | auts2b | pax3a | cyp26b1 | robo4 | hoxa2b | mapk12b | rfx4 | CR751602.2 | wnt7aa | 1 |
| 15 | Ectoderm | Central Nervous System | Neuroblast | Spinal cord | SCProg12a | SC | 1,2 | cdx4 | ldlrad2 | si:ch73-265d | hoxb9a | hoxc8a | hoxb8a | hoxa9a | hoxc6b | ptprz1b | si:ch211-170 | col28a2a | dbx1b | hoxd3a | hoxc1a | wnt4a | hoxb7a | 2 |
| 16 | Mesoderm | Pharyngeal arch | | | Arch2a | Arch2 | 2 | gsc | lhx6 | foxl1 | barx1 | foxf2a | prrx1b | hand2 | pax9 | fgfbp2b | twist1b | fli1a | LO018188.1 | six1a | fibina | phldb2b | prrx1a | 0 |
| 17 | Ectoderm | Central Nervous System | Neuroblast | Anterior spinal cord | antSCProg1a | antSC | 1 | olig3 | gsx2 | olig4 | hoxd4a | zic4 | prss35 | zic1 | zic6 | irx1b | zic5 | rxrga | hoxc1a | fzd10 | msx3 | pax3a | zic2b | 0 |
| 18 | Ectoderm | Neural crest | Cranial neural crest | Cartilage | NCcart2 | NCcart2 | 2 | col9a1a | ccdc3b | fgfbp2b | col11a2 | enpp6 | col9a3 | foxf2a | six2a | cthrc1a | col9a2 | cpxm1a | fibina | foxc1b | angptl1a | cd248a | twist2 | 0 |
| 19 | Ectoderm | Central Nervous System | Neuroblast | Hindbrain | HBProg1c | HB | 1 | egr2b | olig3 | msx1a | zic5 | irx1b | zic4 | msx3 | olig4 | zic1 | fzd10 | si:ch73-364h | zic2b | pax3a | zic2a | rfx4 | CR751602.2 | 0 |
| 20 | Ectoderm | Central Nervous System | Neuron, differentiating | Anterior spinal cord | antSCDiff12a | antSC | 1,2 | hoxd4a | sox21a | gbx1 | rxrga | foxb1a | dbx1b | wfikkn1 | sp8a | plekhh1 | sulf2a | mapk12b | si:ch73-265d | raraa | her8.2 | CR382300.1 | hoxb5b | 0 |
| 21 | Ectoderm | Integument | Integument Gill? | Gill | Gill5b | Gill | 5 | CR626884.5 | cyldb | si:dkey-238k | si:dkey-202l2 | si:dkey-239b | caspbl | plekhs1 | zgc:152791 | si:ch73-329n | AL928650.1 | si:ch211-266 | si:dkey-7c18. | rhcgl1 | si:ch211-226 | si:dkey-201i2 | btr21 | 9 |
| 22 | Mesoderm | Paraxial mesoderm | Somite | Myotome | Myotome | Myotome | 1 | dmrt2a | myf5 | wnt2 | fgl1 | col6a4a | rbp7b | COMP | tcf15 | rprmb | fbln7 | meox1 | cilp | tril | thbs4b | csf1a | mxra5a | 0 |
| 23 | Ectoderm | Integument | Epidermis | Basal cells | Basal5a | BasalEpi5 | 5 | si:dkey-91f15 | ccl27a | col17a1a | htra4 | plek2 | entpd3 | sod3b | cxcl18a.1 | CABZ010442 | mmp30 | ms4a17a.4 | si:ch211-76l2 | si:dkey-95h1 | b3gnt5b | rasgrp4 | cxl34b.11 | 3 |
| 24 | Ectoderm | Neural crest | Cranial neural crest | | NCcranial5a | NCcran5 | 5 | aspn | rgs5a | hpdb | fstl3 | tat | zgc:153704 | CABZ010927 | ttc36 | cldn11a | ifitm1 | abi3bpb | angptl1a | fstb | asip2b | zgc:174904 | ms4a17a.7 | 2 |
| 25 | Ectoderm | Central Nervous System | Neuron | Inhibitory, GABA | FBNeurInhibAll | FB | 1,2,5 | arxb | dlx2b | arxa | dlx1a | gad2 | sp9 | si:dkey-175g | nrxn3a | slc32a1 | gad1b | dlx6a | slc6a1b | nkx2.1 | gria1a | celf4 | gas7a | 1 |
| 26 | Ectoderm | Central Nervous System | Neuron, differentiating | Hindbrain | HBDiff1a | HB | 1 | her3 | dbx1b | dbx1a | robo4 | egr2b | sox21a | gbx1 | wfikkn1 | cyp26b1 | hoxa2b | si:ch73-364h | sp8b | wnt7aa | scube2 | sulf2a | foxb1a | 1 |
| 27 | Ectoderm | Central Nervous System | Neuron | Mid-hindbrain boundary, Glut | MHBNeurGlutAll | MHB | 1,2,5 | si:dkey-10f21 | stmn3 | slc17a6a | zgc:153615 | nos1apb | ppp2r2ca | syn1 | sncgb | si:ch211-113 | adcyap1r1a.1 | grin1a | cacng2a | scn8aa | glra3 | atp2b3a | gphnb | 2 |
| 28 | Mesoderm | Paraxial mesoderm | Parcordal cartilage | | Parachord12 | Parachord | 1,2 | thbs3b | ccdc80l1 | FO704741.1 | ccdc80l2 | si:ch211-160 | apof | alx4a | zgc:162612 | csf1a | si:ch211-69b | cthrc1a | pcolcea | csf1b | ndnfl | col9a1a | tbx15 | 3 |
| 29 | Ectoderm | Neural crest | Cranial neural crest | Pharyngeal arch | NCcranial5b | NCcran5 | 5 | aspn | gsc | CABZ010927 | dkk3b | cldn11a | hpdb | zgc:153704 | rgs5a | ecrg4a | si:dkey-12l12 | angptl1a | abi3bpb | asip2b | fstl3 | mxra8b | ccdc3b | 3 |
| 30 | Ectoderm | Central Nervous System | Neuron | Retinal, differentiating neurons | RetDiff2 | RetDiff | 2 | hes2.2 | six6b | si:dkey-238o | cldn5b | foxn4 | si:ch73-215f | vsx2 | ascl1a | slc38a3a | rx1 | foxg1b | myb | crip3 | nr2e1 | mybl1 | dlc | 2 |
| 31 | Ectoderm | Neural crest | Cranial neural crest | | NCcranial2 | NCcran2 | 2 | spon2b | dlx4a | dlx6a | dlx4b | barx1 | fgfbp2b | dlx1a | dlx2a | dlx5a | foxc1b | grem2b | fli1a | phlda3 | emx2 | dlx3b | fibina | 0 |
| 32 | Ectoderm | Neural crest | Cranial neural crest | | NCcranial1a | NCcran1 | 1 | ednrab | fgf20b | grem2b | barx1 | dlx4b | dlx6a | dlx1a | pdgfbb | dlx2a | phlda3 | cxcr4a | foxc1b | vasna | twist3 | angptl2b | fli1a | 0 |
| 33 | Mesoderm | Paraxial mesoderm | Somite | Sclerotome | Sclero1 | Sclero | 1 | nkx3-1 | aplnr2 | vcanb | pax9 | c7a | rbp7b | COMP | meox1 | tcf15 | angptl7 | mxra5a | twist2 | tril | emilin2b | vwde | cilp | 0 |
| 34 | Ectoderm | Central Nervous System | Neuroblast | Retinal | RetProg1a | RetProg | 1 | rx2 | vsx2 | rx1 | hist1h2a5 | hist1h2a11.4 | zgc:165555 | CR354435.2 | si:dkey-108k | si:dkey-108k | hmx1 | nr2e1 | si:ch1073-15 | si:dkey-261m | si:ch211-113 | hist1h2a2.1 | si:ch1073-15 | 7 |
| 35 | Ectoderm | Central Nervous System | Neuron, differentiating | Ciliated, spinal cord | SCNeurCilia12a | SC | 1,2 | rsph9 | cfap52 | spag6 | tbata | cfap126 | si:ch211-153 | armc4 | si:ch211-163 | si:dkeyp-110 | pih1d2 | spata18 | si:ch211-163 | cdx4 | lrrc6 | si:dkey-148f1 | lrrc23 | 5 |
| 36 | Mesoderm | Muscle | Skeletal muscle | Prog? | MusSkelProgAll | MusSkelProg | 1,2,5 | mymk | filip1a | flncb | cdh15 | klhl40b | asb12b | BX649294.1 | chrng | CR925737.1 | trim109 | FO818704.2 | mss51 | alpk3b | AL772146.2 | klhl40a | tmx2a | 0 |
| 37 | Ectoderm | Central Nervous System | Neuron | Retinal, differentiating neurons | RetDiff25a | RetDiff | 2,5 | atoh7 | tfap2d | prdm1b | ndrg1b | hes2.2 | foxn4 | mybl1 | otx2b | slc38a3a | foxg1b | cspg5b | ptprua | bhlhe22 | tbx2b | six6b | neurod4 | 0 |
| 38 | Mesoderm | Blood | Hematopoietic cell | Nucleate erythrocyte | RBC1a | RBC1 | 1 | CR854926.1 | tfr1a | drl | CR388184.1 | susd1 | gfi1b | slc10a4 | fech | si:dkey-261j4 | si:dkey-150i1 | gpr78a | tbx16l | cyp46a1.1 | nfe2 | gata1a | ppox | 2 |
| 39 | Ectoderm | Central Nervous System | Neuroblast | Spinal cord | SCProg12b | SC | 1,2 | lbx1b | hoxc8a | hoxc9a | hoxb6b | prdm13 | hoxc6b | ldlrad2 | wnt4a | hoxa9a | hoxb9a | hoxb8a | si:ch211-67e | si:dkeyp-110 | prss35 | hoxc1a | olig3 | 2 |
| 40 | Ectoderm | Central Nervous System | Neuroblast | Fore-midbrain boundary | FMBProgAll | FMB | 1,2,5 | si:ch211-69b | zic6 | wnt8b | tpbga | otx1 | otx2a | rspo2 | cep112 | barhl2 | zic4 | zic5 | wnt4a | sp5a | tnfrsf19 | clybl | otx2b | 1 |
| 41 | Ectoderm | Central Nervous System | Neuroblast | Mid-hindbrain boundary | MHBProgAll | MHB | 1,2,5 | pax5 | en1a | en2a | en1b | fgf8a | slf1 | gbx2 | atoh1a | pax2a | en2b | sp9 | oc90 | olig3 | tubb2 | ism1 | il17rd | 0 |
| 42 | Ectoderm | Central Nervous System | Neuron, differentiating | Midbrain, cerebellum | MBDiff25 | MB | 2,5 | helt | zgc:101100 | hs3st1l2 | CU929451.2 | masp1 | CR848047.1 | ptprz1a | gsx1 | si:ch73-215f | otx2b | pax7a | ascl1b.1 | cep112 | nfia | znf219 | nr2e1 | 2 |
| 43 | Ectoderm | Central Nervous System | Neuroblast | Hindbrain, ventral | HBProg1d | HB | 1 | mpz | egr2b | nkx6.2 | gbx1 | sox21a | cyp26b1 | nkx6.1 | sulf2a | hoxa2b | wfikkn1 | foxb1a | CR848812.1 | prdm12b | sulf2b | prdm8 | sfrp1a | 0 |
| 44 | Mesoderm | Paraxial mesoderm | Somite | Myoblast | MyoProg5a | MyoProg | 5 | msc | col18a1b | myf5 | myog | myod1 | txnipb | jam2a | vwde | jam2b | chrna1 | zgc:92518 | cyp2n13 | pitx3 | hsp90aa1.1 | mafaa | tcf15 | 1 |
| 45 | Endoderm | Pharyngeal endoderm | Pharyngeal epidermis | | PharEpi1a | PharEpi1 | 1,2 | pax1b | si:dkey-43p1 | pax1a | nkx2.7 | fgf17 | foxi1 | nkx2.3 | frem2a | shhb | foxa2 | fgf24 | irx7 | tbx1 | ptprfa | sostdc1a | fgfrl1b | 1 |
| 46 | Endoderm | Pharyngeal endoderm | Pharyngeal epidermis | | PharEpi1b | PharEpi1 | 1,2 | foxi1 | vgll4l | padi2 | si:ch211-67e | ppp1r1b | kera | fermt1 | lrata | cldn1 | col7a1 | zgc:171775 | col14a1a | zgc:101810 | sostdc1a | ecrg4b | si:ch211-202 | 4 |
| 47 | Ectoderm | Integument | Epidermis | Basal cells | Basal2a | BasalEpi2 | 2 | si:ch211-256 | si:dkey-95h1 | col28a1a | col17a1a | CU459012.1 | zgc:100997 | si:ch211-241 | col17a1b | lamb4 | entpd3 | si:ch211-213 | col5a3b | lrata | zgc:113223 | c7a | si:ch211-76l2 | 6 |
| 48 | Mesoderm | Blood | Hematopoietic cell | Nucleate erythrocyte | RBC2b | RBC2 | 2 | BX005417.1 | si:ch73-288o | si:ch211-250 | gp9 | hsp70l | CR293511.1 | si:ch211-207 | zgc:163057 | nmt1b | cabp2a | hbae5 | epb41b | fbxo32 | slc20a1a | alas2 | nt5c2l1 | 4 |
| 49 | Nonspecific genes | | | | Nonspecific12 | Nonspecific | 1,2 | crip3 | six3b | prmt3 | umps | fabp11a | myca | tjp2b | rnaseh2b | kmt5ab | chchd10 | cmss1 | kpna2 | hgh1 | atic | mcmbp | aprt | 0 |
| 50 | Ectoderm | Neural crest | Cranial neural crest | Cartilage | NCcart5a | NCcart5 | 5 | otos | tnmd | angptl5 | ucmab | epyc | adgrg6 | cpxm1a | col9a1a | col11a2 | aspn | angptl1a | fstl3 | zgc:153704 | zgc:113307 | pigr | col9a3 | 2 |
| 51 | Ectoderm | Central Nervous System | Neuroblast | Hindbrain? | HBDiff12a | HB | 1,2 | AL845362.1 | lbx1b | prdm13 | gsx2 | tfap2b | BX530077.1 | kirrel3l | ascl1a | pax3a | zic1 | draxin | pcdh1g32 | zic4 | hoxd4a | CR848812.1 | zic2b | 0 |
| 52 | Ectoderm | Central Nervous System | Neuroblast | Retinal | RetProg0a | RetProg | 1 | vsx2 | rx1 | si:dkey-238o | hmx4 | six3b | crip3 | si:dkey-261m | tjp2b | fabp11a | umps | si:dkey-108k | exosc8 | pax6b | rnaseh2b | trmt6 | myca | 3 |
| 53 | Mesoderm | Paraxial mesoderm | Somite | Myoblast | MyoProg5b | MyoProg | 5 | si:ch211-287 | clec3ba | fgl1 | cldn7a | olfml2bb | myf5 | admb | dmrt2a | ackr3a | fbln7 | rbp7b | zgc:113307 | thbs4b | cilp | si:ch211-243 | csf1b | 3 |
| 54 | Ectoderm | Central Nervous System | Neuroblast | Retinal | RetProg0b | RetProg | 1 | rx1 | six3b | foxd1 | fabp11a | fahd1 | umps | cx43 | myca | prmt3 | cmss1 | zgc:194224 | dctpp1 | ftsj3 | exosc8 | ddx51 | atic | 1 |
| 55 | Endoderm | Liver | Hepatocyte | | Hepato5a | Liver | 5 | cbln14 | si:ch211-113 | zgc:171534 | or109-11 | serpina7 | cbln5 | cbln9 | ccl39.2 | zgc:172253 | cpb2 | ahsg1 | si:ch211-262 | si:ch73-15n2 | si:dkey-96f10 | itih2 | si:dkeyp-73d | 7 |
| 56 | Ectoderm | Neuron | Cranial ganglia | | CranGangAlla | CranGang | 1,2,5 | pvalb6 | zgc:158291 | tlx2 | cerkl | isl2b | rab3db | gfra1a | six2b | ngfrb | six4b | stk25a | phox2a | isl2a | msi2a | neurod1 | eya2 | 1 |
| 57 | Mesoderm | Muscle | Skeletal muscle | Slow muscle cell | MusSkelSlow1a | MusSkelSlow | 1 | si:dkeyp-68b | CU896691.1 | kcnmb2 | si:ch211-28p | CU633479.2 | smyhc3 | klhl41a | actn2b | smyhc2 | rnf207b | myh7bb | mylk3 | BX088524.3 | myom1b | tnni1d | ryr1a | 2 |
| 58 | Ectoderm | Central Nervous System | Neuron | Retinal, differentiating neurons | RetDiffAll | RetDiff | 1,2,5 | atoh7 | mybl1 | hes2.2 | foxn4 | dlc | bhlhe22 | six6b | foxg1b | pou2f2a.1 | rx1 | vsx1 | onecut1 | dhx32b | tbx2b | insm1a | neurod4 | 0 |
| 59 | Endoderm | Thymus | | | Thymus | Thymus | 1,2,5 | cytip | si:ch211-214 | si:ch73-208g | sla2 | ccl38.6 | lpar5a | CU019662.1 | vaspa | si:dkeyp-68b | si:dkey-185m | prkacba | rasal3 | ccr9a | wipf1a | wasa | tagapb | 4 |
| 60 | Ectoderm | Central Nervous System | Neuron | Retinal, differentiating neurons | RetDiff25b | RetDiff | 2,5 | nr2e3 | samd7 | six7 | otx5 | prdm1b | crx | tmx3a | zgc:109965 | si:ch73-28h2 | gngt2a | tulp1a | pde6gb | lhx4 | fkbp1ab | elovl4b | ndrg1b | 2 |
| 61 | Ectoderm | Central Nervous System | Neuroblast | HB,MN | HBProgMN1e | HB | 1 | nkx2.9 | nkx2.2b | nkx6.2 | nkx2.2a | foxa | ntn1b | helt | phox2bb | foxa2 | nkx6.1 | si:ch73-265d | sulf2b | si:ch73-364h | ntn1a | npas1 | wnt7aa | 2 |
| 62 | Ectoderm | Central Nervous System | Neuroblast | Forebrain | FBProgAllb | FB | 1,2,5 | rx3 | nkx2.4b | nkx2.4a | nkx2.1 | fzd8b | six6a | shha | ndnfl | si:dkeyp-110 | vax1 | sox1b | fezf1 | six3a | fezf2 | nrp2b | nr2e1 | 1 |
| 63 | Mesoderm | Blood | Hematopoietic cell | Nucleate erythrocyte | RBC1b | RBC1 | 1 | drl | tfr1a | si:dkey-261j4 | CR388184.1 | gfi1b | CU463284.1 | susd1 | si:dkey-150i1 | slc10a4 | btk | slc25a37 | ppox | cyp46a1.1 | gata1a | cldng | slc9a1b | 2 |
| 64 | Ectoderm | Neural crest | Neural Progenitors | | NCneural12 | NCneural | 1,2 | erbb3b | crestin | mov10b.1 | mmp17b | ednrab | sox10 | foxd3 | mcamb | BX248318.1 | prdm12b | ngfrb | slc15a2 | mcf2l2 | snai1b | CABZ010312 | myo10l3 | 0 |
| 65 | Ectoderm | Central Nervous System | Neuron, differentiating | Anterior spinal cord | antSCDiff12b | antSC | 1,2 | si:ch211-152 | BX530077.1 | lbx2 | her13 | prdm8 | nkx6.1 | cnksr2b | sertad2a | prox1a | onecutl | gadd45gb.1 | btbd6a | neurog1 | LIN28A | pcdh1g32 | inavaa | 1 |
| 66 | Ectoderm | Central Nervous System | Neuron, differentiating | Hindbrain | HBDiff12b | HB | 1,2 | BX530077.1 | dbx1a | dbx1b | otpb | onecutl | neurog1 | her13 | gadd45gb.1 | inavaa | ascl1b.1 | nhlh2 | prox1a | neurod4 | sp8b | srrm4 | cbfa2t2 | 0 |
| 67 | Ectoderm | Central Nervous System | Floor plate | Midbrain | FloorplateMB1 | Floorplate | 1 | foxa2 | foxa | shha | foxa1 | shhb | ntn1b | ldlrad2 | nkx6.2 | pitx3 | spon1b | otx2a | lmx1bb | ntn1a | npas1 | ascl1b.1 | otx1 | 0 |
| 68 | Ectoderm | Central Nervous System | Neuron, differentiating | Fore-midbrain boundary | FMBDiff12a | FMB | 1,2 | emx1 | tbr1b | srgap1b | myt1la | mpped1 | cpne2 | eomesa | kctd4 | kcnip1b | l1cama | glipr1b | PRKAR2B | neurod6b | khdrbs2 | olfm1a | etv1 | 0 |
| 69 | Ectoderm | Neural crest | Cardiac neural crest | | NCcardiac | NCcard | 1 | zgc:162612 | csf1a | FO704741.1 | alx4a | ccdc80l2 | arhgap36 | emilin1a | npb | csf1b | cldn11a | rasl11a | foxc1a | cthrc1a | enpp6 | si:ch211-267 | jcada | 2 |
| 70 | Ectoderm | Integument | Epidermis | Basal cells | Basal1 | BasalEpi1 | 1 | itgb3a | si:ch211-218 | si:dkey-117n | col7a1l | thbs3a | si:dkey-117n | col5a3b | cpz | CU469568.1 | def6c | lamb4 | olfm2b | ptger1a | si:ch211-256 | si:dkey-91f15 | lrata | 5 |
| 71 | Mesoderm | Myeloid lineage | Monocyte | Macrophage | Macro5 | Macro | 5 | itgae.1 | c1qa | NPC2 (1 of m | si:dkey-148a | zgc:103700 | marco | c1qb | mrc1b | ccl35.1 | si:ch211-194 | BX682234.1 | csf2rb | si:dkey-1h24. | slc2a6 | ccl34a.4 | CABZ010881 | 4 |
| 72 | Ectoderm | Central Nervous System | Neuron, differentiating | Mid-hindbrain boundary | MHBDiff25 | MHB | 2,5 | scrt1b | tfap2d | sox12 | insm1b | zgc:162707 | stap2a | scrt2 | mpp2a | en2b | lef1 | insm1a | lhx9 | ptprua | en2a | lzts1 | srrm4 | 1 |
| 73 | Ectoderm | Central Nervous System | Neuron | Retinal, mix of RGC and inhibitory? (a | RetNeuron25 | RetNeuron | 2,5 | pax10 | si:ch73-290k | prdm13 | gad2 | tfap2b | sox12 | barhl2 | slc32a1 | bhlhe22 | six6b | foxg1b | lhx9 | ndrg4 | kctd15a | scrt1a | pde6gb | 1 |
| 74 | Ectoderm | Central Nervous System | Neuroblast | Retinal Muller Glia? RPE? | RetProgAllb | RetProg | 1,2,5 | vsx2 | rx1 | hmx1 | col15a1b | si:dkey-238o | nr2e1 | hmx4 | six3b | myb | six3a | wfdc1 | crip3 | ggctb | cart3 | dclk2a | tbx2a | 1 |
| 75 | Ectoderm | Central Nervous System | Neuron, differentiating | Hindbrain | HBDiff25 | HB | 2,5 | masp1 | CU929451.2 | AL845362.1 | CR848047.1 | atoh1a | nfia | si:ch73-215f | olig3 | znf219 | zic4 | zic5 | CU467822.1 | chfr | zic1 | prss35 | zic2a | 1 |
| 76 | Ectoderm | Central Nervous System | Neuron | Mid-hindbrain boundary, Inhibitory, G | MHBNeurGABAAll | MHB | 1,2,5 | sprn | aig1 | sv2a | nell2b | pcsk1nl | scg2b | eno2 | sypa | gpr85 | kif5aa | scg5 | si:dkey-12h9. | itm2ca | lhx1a | maptb | atp1a3a | 1 |
| 77 | Ectoderm | Integument | Periderm | | Peri2a | Peri2 | 2 | ponzr4 | BX571705.1 | si:ch211-161 | ubap1la | wu:fj05g07 | zgc:112964 | trim35-9 | grk5 | b3gnt3.1 | cyp17a1 | si:rp71-1h20. | ftr86 | si:ch211-264 | jac10 | si:ch73-196i1 | si:dkeyp-13a | 6 |
| 78 | Ectoderm | Integument | Periderm | | Peri2b | Peri2 | 2 | CABZ010440 | CABZ010440 | CABZ010676 | si:ch1073-19 | si:ch1073-34 | si:dkey-73p2. | jac10 | si:dkey-73p2. | si:ch73-196i1 | CABZ010720 | CABZ010155 | zgc:112964 | ponzr3 | si:dkey-239b | nos1 | zgc:171566 | 8 |
| 79 | Mesoderm | Blood vessel | Endothelial | | EndotheAlla | Endothelial | 1,2,5 | mrc1a | lyve1b | sele | stab2 | CU927890.1 | gpr182 | rasip1 | stab1 | tie1 | myct1a | tek | scarf1 | si:ch211-33e | myct1b | flt4 | clec14a | 1 |
| 80 | Ectoderm | Neural crest | Cranial neural crest | | NCcranial5e | NCcran5 | 5 | cyp1c1 | slco2b1 | cldn11a | zgc:153704 | timp4.3 | agtr2 | CABZ010927 | cyp2ad2 | fstb | serpinf1 | hpdb | fstl3 | zgc:162612 | cyp2p7 | zgc:174904 | hgd | 3 |
| 81 | Ectoderm | Integument | Epidermis | Fin basal cells | BasalFin5a | BasalEpiFin5 | 5 | si:ch211-251 | cxl34b.11 | and3 | epgn | postna | ms4a17a.5 | ms4a17a.1 | plek2 | b3gnt5b | mmp30 | rab42b | si:dkey-95h1 | si:dkey-102c | cxcl8a | zgc:113223 | cldn1 | 4 |
| 82 | Ectoderm | Central Nervous System | Neuron, differentiating | Hindbrain | HBDiff12c | HB | 1,2 | atoh1b | si:ch211-215 | scrt1b | onecutl | scrt1a | lzts1 | nrn1a | nkain2 | sdk1a | gdf11 | lhx9 | uncx | FP017274.1 | nhlh2 | pdlim5a | mpp2a | 1 |
| 83 | Ectoderm | Central Nervous System | Neuron | Retinal, differentiating neurons, epipl | RetDiff25c | RetDiff | 2,5 | zgc:112294 | zgc:103625 | rpgrip1 | slc1a8a | opn6a | foxq2 | CABZ011176 | glmnb | arhgap20 | si:ch211-276 | zgc:109965 | tmem136b | rcvrna | sez6a | samd7 | BX957322.3 | 4 |
| 84 | Mesoderm | Paraxial mesoderm | Somite | Myoblast | MyoProg2a | MyoProg | 2 | myog | myod1 | myf5 | jam2b | tbx15 | jam2a | pitx3 | efhd1 | hsp90aa1.1 | chrna1 | pitx2 | klhl41b | tbx1 | zgc:92518 | unc45b | stac3 | 1 |
| 85 | Mesoderm | Blood | Hematopoietic cell | Nucleate erythrocyte | RBC2c | RBC2 | 2 | gp9 | hif1al2 | slc9a1b | klf1 | BX284666.1 | tbx16l | si:ch73-250a | si:dkey-261j4 | gata1a | CR293511.1 | rfesd | si:dkey-93h2 | si:ch211-250 | rell2 | fech | epb41b | 4 |
| 86 | Ectoderm | Central Nervous System | Neuron, differentiating | Mid-hindbrain boundary | MHBDiff12b | MHB | 1,2 | her5 | her11 | pax5 | pax2b | en2a | irx7 | zgc:101100 | en1b | en2b | rtn4r | pax2a | nkx6.2 | BX571942.1 | otx1 | foxb1a | otx2a | 1 |
| 87 | Ectoderm | Neural crest | Cranial neural crest | Arch mesenchyme from crest | NCcranial1b | NCcran1 | 1,2 | nkx3.3 | foxf1 | tmem98 | hlx1 | foxf2b | foxf2a | isl2b | BX088707.3 | nkx2.3 | gfra1a | osr1 | rbpms2a | rbpms2b | barx1 | isl2a | inhbaa | 0 |
| 88 | Mesoderm | Blood vessel | Endothelial | | EndotheAllb | Endothelial | 1,2,5 | notchl | myct1b | flt1 | pecam1 | si:ch211-33e | tie1 | ecscr | myct1a | scarf1 | rasip1 | ccm2l | ushbp1 | micall2a | arhgef9b | clec14a | f2rl2 | 1 |
| 89 | Ectoderm | Central Nervous System | Neuroblast | Forebrain | FBProgAllc | FB | 1,2,5 | nkx2.4a | nkx2.4b | nkx2.1 | fezf2 | emx2 | sp5a | emx3 | sox1b | lhx2b | pcdh10a | oc90 | nrp2b | efhd1 | nr2e1 | lhx5 | foxd2 | 0 |
| 90 | Ectoderm | Central Nervous System | Oligodendrocyte | Forebrain | OligoFB | Oligo | 1,2,5 | olig1 | nkx2.2a | nkx2.2b | nkx2.9 | gsx2 | vax1 | olig2 | lhx5 | foxd1 | ntn1b | sox1b | robo4 | wnt7bb | nr2e1 | npas1 | podxl | 0 |
| 91 | Ectoderm | Central Nervous System | Neuroblast | Forebrain | FBProgAlld | FB | 1,2,5 | eno4 | vax1 | fezf1 | sox1b | fgf8a | six6a | ntn1b | fezf2 | cyp26c1 | otpb | six3a | pax2a | lhx2b | nr2e1 | pdgfaa | zic5 | 0 |
| 92 | Mesoderm | Paraxial mesoderm | Somite | Sclerotome | Sclero2 | Sclero | 2 | aplnr2 | c7a | pax9 | emilin2b | agtr2 | cpxm1a | nkx3.2 | kazald3 | zgc:154093 | wfdc2 | zgc:162612 | col5a2a | fgfrl1b | ednraa | twist1b | aebp1 | 2 |
| 93 | Ectoderm | Placode | Otic | | Otic | Otic | 1,2,5 | otomp | bricd5 | si:ch211-80h | otog | cldna | si:dkey-30e9. | si:dkey-16l2. | gal3st1a | si:dkey-222f2 | si:ch211-137 | zgc:112437 | atp1a1a.4 | stm | mb | si:ch211-243 | si:ch1073-18 | 8 |
| 94 | Ectoderm | Placode | Neuron | Olfactory placode | Olfactory | Olfactory | 1,2,5 | ompb | lhx2a | si:ch73-335l2 | abhd2b | cnga4 | rasd3 | got1l1 | vsig8a | ppp1r15b | fabp10b | b3gat1b | capslb | mef2b | fam228a | emb | cntn4 | 1 |
| 95 | Ectoderm | Central Nervous System | Neuron, differentiating | Spinal cord | SCDiff12a | SC | 1,2 | chrna2a | nkx1.2lb | kcnd3 | si:ch211-215 | si:ch211-237 | sall3a | plekho1a | zgc:77784 | CR974456.1 | onecut2 | scube1 | onecutl | sv2bb | LIN28A | sox14 | gata2a | 2 |
| 96 | Ectoderm | Retina | Epithelial | Retinal pigmented epithelium | RPE | RPE | 5 | cracr2ab | si:dkey-43k4. | pmelb | msnb | slc24a5 | rlbp1a | si:dkeyp-86d | tyrp1a | sytl2b | mitfa | rlbp1b | slc45a2 | fabp11b | dct | FP085398.1 | cax1 | 2 |
| 97 | Ectoderm | Central Nervous System | Neuroblast | Radial glia, hindbrain | RadialGlia2 | RadGlia | 2 | CU929451.2 | hepacama | NPAS3 | masp1 | slc6a1b | CR848047.1 | gpsm1a | si:ch1073-30 | si:ch73-215f | npas1 | nfia | chfr | ptn | her8.2 | ptprz1b | lrrtm2 | 2 |
| 98 | Mesoderm | Pectoral fin bud | | | FinBud12 | FinBud | 1,2 | tbx5a | tbx18 | csf1b | hoxa13b | bmp2a | tbx15 | hand2 | fgf10a | prrx1b | hoxd11a | hoxd12a | twist3 | hoxd9a | alx4a | hoxd10a | notum1b | 0 |
| 99 | Ectoderm | Central Nervous System | Neuroblast | Retinal Muller Glia? RPE? | RetProgAllc | RetProg | 1,2,5 | aldh1a3 | vax1 | bmp7b | c1qtnf12 | rx1 | cyp27c1 | vsx2 | fezf2 | stra6 | col15a1b | caska | six3b | cx43 | pou2f2a | lhx2b | pdzrn3b | 0 |

| | | | | | | | | | | | | | | | | | | | | | | | | |
|---|---|---|---|---|---|---|---|---|---|---|---|---|---|---|---|---|---|---|---|---|---|---|---|---|
| 100 | Mesoderm | Mesenchyme-related, arch or fin bud? | | | MesenUnk2 | MesenUnk2 | 2 | osr2 | mkxa | alx4a | enpp6 | zgc:113307 | pitx2 | adma | FQ323119.1 | cthrc1a | lum | calhm2 | rhoq | cpxm1a | pitx3 | cldn11a | cd248a | 1 |
| 101 | Endoderm | Intestine | Intesinal epithelial cell | | IntestineEpiAll | IntestineEpi | 1,2,5 | ihha | pdx1 | hnf1ba | hnf4g | cldnc | gata6 | cldn15a | zgc:193726 | slc13a2 | si:ch211-248 | nr5a2 | foxa3 | aqp12 | hnf4a | eps8l3a | sst1.2 | 2 |
| 102 | Ectoderm | Neural crest | Cranial neural crest | | NCcranial1c | NCcran1 | 1 | ednrab | grem2b | si:dkey-68o6. | dlx2a | prrx1b | twist1a | phlda3 | prrx1a | foxc1b | snai1a | tuba8l3 | snai2 | pmp22a | rbms2b | stxbp6 | tpm4a | 1 |
| 103 | Mesoderm | Muscle | Skeletal muscle | Fast muscle cell | MusSkelFast2b | MusSkelFast | 2 | calcoco1b | igfn1.3 | mybpha | ankrd1b | CABZ010707 | zgc:136908 | myha | prkg1b | hsc70 | ampd1 | hspb6 | pfkma | CU693479.1 | tmem41ab | atp2a1l | tnni2b.2 | 1 |
| 104 | Ectoderm | Central Nervous System | Neuron | Retinal, differentiating neurons | RetDiff25d | RetDiff | 2,5 | nr2e3 | samd7 | BX957322.3 | prdm1b | six7 | CABZ010555 | otx5 | tulp1a | sept4a | tmx3a | ndrg1b | zgc:109965 | crabp1a | dscamb | crx | si:dkey-238o | 2 |
| 105 | Ectoderm | Neuron | Cranial ganglia | | CranGangAllb | CranGang | 1,2,5 | sst6 | tppp2 | emb | tmeff2b | drgx | ppp1r1c | tlx2 | pou4f1 | zgc:158291 | ank2a | CABZ011187 | asphd1 | foxg1d | stmn3 | six2b | atp2b3b | 1 |
| 106 | Ectoderm | Lens placode | Lens epithelium | | LensEpi | Lens | 1,2,5 | foxe3 | CR356246.1 | gja8b | slc7a11 | prox2 | FO681288.1 | col28a1a | vcana | cx23 | sparcl1 | lim2.1 | pitx3 | col4a3 | pltp | mafb | hmx1 | 0 |
| 107 | Ectoderm | Central Nervous System | Neuron | Retinal, differentiating neurons | RetDiff2 | RetDiff | 2 | CABZ010555 | sept4a | si:dkey-238o | crabp1a | vsx1 | cldn5b | cspg5b | six6b | hes2.2 | ptprua | olig2 | si:ch73-215f7 | CR848047.1 | foxn4 | foxg1b | sh3bgrl2 | 2 |
| 108 | Ectoderm | Integument | Epidermis | Fin basal cells | BasalFin1 | BasalEpiFin1 | 1 | si:ch1073-80 | itgb3a | msx2a | col4a4 | CU469568.1 | cpz | col5a3b | lamb4 | si:ch211-241 | zgc:154006 | fgfbp1b | olfm2b | si:dkey-91f15 | myof | col4a3 | wnt6b | 4 |
| 109 | Ectoderm | Central Nervous System | Neuroblast | Radial glia, mid-hindbrain | RadialGlia5 | RadGlia | 5 | atoh1c | BX469902.1 | si:ch211-66e | hepacama | mfge8a | IGLON5 | ptn | zbtb4 | eno1b | clybl | nfia | zgc:194659 | cspg5b | slc6a1b | slc4a4a | crtac1a | 2 |
| 110 | Mesoderm | Pectoral fin bud | | | FinBud25 | FinBud | 2,5 | hoxb13a | si:dkey-202e | c1qtnf5 | hoxc13b | hoxc13a | hoxa13a | hoxa13b | CABZ010741 | bmp2a | and1 | angptl5 | angptl7 | mxra8b | kazald3 | ttc36 | efemp2b | 1 |
| 111 | Ectoderm | Central Nervous System | Neuroblast | Midbrain | MBProg12b | MB | 1,2 | dmbx1a | zic6 | si:ch211-69b | cep112 | hs3st1l2 | hey2 | tnfrsf19 | fzd10 | BX571942.1 | pax7b | otx1 | pax7a | msx1a | zic4 | en2b | irx1a | 1 |
| 112 | Ectoderm | Central Nervous System | Neuron | Spinal cord glycinergic | SCNeurInhib12a | SC | 1,2 | lbx1a | skor1a | zgc:77784 | si:dkey-175g | pax2b | slc6a5 | scrt1a | nkain2 | ssx2ipb | si:ch211-215 | plekho1a | pdlim5a | zgc:165603 | hmx3a | lhx1a | onecutl | 4 |
| 113 | Ectoderm | Central Nervous System | Neuron | Retinal, differentiating neurons | RetDiff25e | RetDiff | 2,5 | zgc:158291 | pou6f2 | irx4a | pou4f2 | barhl1a | si:ch73-290k | pax10 | atoh7 | tfap2d | rbpms2b | inab | kif3cb | isl2b | gng13b | isl1 | pou4f1 | 2 |
| 114 | Ectoderm | Central Nervous System | Neuroblast | Anterior spinal cord | antSCProg1b | antSC | 1 | lbx1b | gsx1 | hoxd4a | pax7a | angptl2b | wnt7bb | hoxb5b | pax7b | pax3a | sox21a | plekhg7 | prdm8 | crabp1b | sp8a | foxb1a | hoxc1a | 0 |
| 115 | Ectoderm | Retina | Photoreceptor | | RetPR | RetPR | 5 | rhol | cnga1a | rom1a | ndufs8b | CT573282.1 | cabp4 | si:rp71-39b2 | gucy2f | si:dkeyp-57d | slc1a2a | cngb3.1 | cnga3b | zgc:77752 | si:ch73-352p | abca4b | grk1a | 4 |
| 116 | Mesoderm | Muscle | Skeletal muscle | Slow muscle cell | MusSkelSlow25 | MusSkelSlow | 2,5 | tnnt2e | si:ch211-89o | CU633479.5 | zgc:195001 | tnni1d | smyhc3 | CABZ010766 | CU633479.2 | rnf207b | map3k7cl | ryr1a | mybpc3 | mybpc1 | smyhc2 | hhatlb | myh7bb | 2 |
| 117 | Mesoderm | Lateral plate mesoderm | VesselProg | | Vessel1 | Vessel1 | 1 | tcf21 | esm1 | pitx3 | tbx1 | gfra3 | si:ch211-160 | tmem88b | pdgfrb | aplnra | foxf2a | pdgfab | si:dkey-261h | nrp2a | jcada | adam8a | ngfb | 2 |
| 118 | Ectoderm | Integument | Epidermis | Basal cells | Basal12 | BasalEpi1 | 1,2 | si:dkey-18p1 | itgb3a | tlx1 | arhgef25b | dkk1a | def6c | cyp26a1 | lrata | adra1d | col4a4 | si:ch211-213 | col4a3 | wnt6b | adam28 | nkx2.7 | nkx2.3 | 2 |
| 119 | Mesoderm | Hypoblast | Hatching gland | | HG1 | Hatching | 1 | si:dkey-239j1 | si:dkey-269i1 | si:dkey-26g8. | zgc:174154 | zgc:174153 | he1.2 | si:dkey-269i1 | si:dkey-239j1 | ctslb | he1.1 | zgc:174855 | si:dkey-108k | fabp11a | exosc8 | rrm2 | cmss1 | 9 |
| 120 | Ectoderm | Central Nervous System | Neuroblast | Hindbrain | HBProg12 | HB | 1,2 | zgc:77784 | lhx1a | pax2b | scrt1a | otpb | FP017274.1 | si:ch211-215 | onecutl | pax2a | plekho1a | mpp2a | lzts1 | otpa | nkain2 | zgc:162707 | lhx5 | 3 |
| 121 | Endoderm | Liver | Hepatocyte | | Hepato5b | Liver | 5 | si:ch211-186 | cbln17 | cyp2x10.2 | tdrd5 | apobb.2 | AL953907.1 | crp2 | igfals | zgc:103559 | cfhl3 | uroc1 | gys2 | f9b | acsbg1 | CR626907.1 | fgf19 | 2 |
| 122 | Ectoderm | Central Nervous System | Neuroblast | Retinal Muller Glia? | RetProg5a | RetProg | 5 | slc38a3a | crabp1a | cldn5b | rx1 | foxn4 | foxg1b | si:dkey-238o | six6b | crip3 | ascl1a | myb | wfdc1 | aplnra | enc3 | cx43 | si:ch73-215f7 | 2 |
| 123 | Ectoderm | Central Nervous System | Neuron, differentiating | Spinal cord | SCDiff2 | SC | 2 | CU929451.2 | si:dkeyp-110 | hoxc9a | lrrc4.1 | hoxa9b | npas1 | zgc:113531 | ldlrad2 | hoxa9a | hoxb8a | hoxb10a | si:ch73-265d | hoxb9a | hoxd9a | CR848047.1 | hoxc8a | 3 |
| 124 | Ectoderm | Central Nervous System | Neuron, differentiating | Fore-midbrain boundary | FMBDiff12b | FMB | 1,2 | eomesa | tbr1b | emx3 | neurod6b | foxg1a | lhx2b | lhx9 | fezf1 | adam22 | fezf2 | bcl11ba | uncx4.1 | nrn1a | znf296 | nfia | clvs2 | 2 |
| 125 | Mesoderm | Pectoral fin bud | Apical epidermal ridge? | | FinBud5 | FinBud | 5 | c1qtnf5 | si:ch211-14k | hoxa13b | hoxa13a | mxra8a | adgrg6 | chst2b | loxa | si:ch211-76l2 | bgnb | notum1b | CABZ010927 | tat | pah | timp4.3 | plcd4b | 2 |
| 126 | Ectoderm | Central Nervous System | Neuron, differentiating | HB,MN | HBDiffMN12d | HB | 1,2 | tbx20 | CR812832.1 | phox2a | tm6sf2 | phox2bb | chodl | rap1gap | prph | inab | pdlim5a | onecutl | isl1 | si:ch211-215 | nkx6.1 | tp53bp2b | dachb | 1 |
| 127 | Mesoderm | Paraxial mesoderm | Somite | Myoblast | MyoProg2b | MyoProg | 2 | fgl1 | si:ch211-287 | fbln7 | zgc:113307 | loxa | fap | pdgfrl | dmrt2a | en1a | pcp4l1 | spon2b | col6a2 | col6a3 | si:ch1073-45 | stk17a | meox1 | 3 |
| 128 | Ectoderm | Integument | Integument Ionocyte | NaK Ionocyte | IonoNaK | IonoNaK | 1,2,5 | kcnj1a.3 | kcnj1a.6 | kcnj1a.2 | bmp8a | foxi3b | foxi3a | sstr5 | atp1b1b | atp1a1a.2 | agrp | CR356236.1 | si:ch73-14h1 | cers3a | mal2 | prg4a | BX000438.2 | 1 |
| 129 | Ectoderm | Neural crest | Iridiphore | | NCirid | NCirid | 1,2,5 | unm_sa821 | CR318624.2 | si:ch73-89b1 | si:ch211-38m | si:ch211-256 | akap12a | hsf5 | fhl2b | alx4b | zgc:110789 | emp3b | si:dkey-197i2 | apoda.1 | gpnmb | tcirg1a | gpx2 | 5 |
| 130 | Mesoderm | Lateral plate mesoderm | Heart primordium | Heart tube | HeartTube | Heart | 1,2 | angptl6 | gata5 | tmem98 | gata6 | clec19a | jam2b | pitx1 | tmem88a | hand2 | jcada | sema3e | bmp6 | pcolcea | FQ323119.1 | col6a3 | BX005003.1 | 0 |
| 131 | Ectoderm | Central Nervous System | Neuron, differentiating | Hindbrain | HBDIff1b | HB | 1 | tm6sf2 | CR812832.1 | nkx2.2b | phox2bb | phox2a | nkx2.9 | nkx2.2a | nkx6.2 | foxa | nkx6.1 | ntn1b | spon1b | fam117ab | si:ch73-265d | CR847953.1 | ascl1b.1 | 1 |
| 132 | Ectoderm | Central Nervous System | Oligodendrocyte | Fore-midbrain boundary | OligoFMB | Oligo | 1,2,5 | nkx2.4a | nkx2.4b | wnt16 | olig1 | wnt8b | nkx2.1 | dbx1b | shha | lhx5 | pitx2 | ntn1b | barhl2 | fezf2 | foxb1a | vegfab | olig2 | 0 |
| 133 | Ectoderm | Integument | Integument Ionocyte | NCC Ionocyte | IonoNCC5a | IonoNCC | 5 | si:dkey-189h | slc12a10.2 | clcn2c | kcnj1a.1 | atp1a1a.3 | si:ch73-14h1 | si:ch73-359m | atp1a1a.2 | foxi3b | prlra | CABZ010834 | BX000438.2 | prom2 | si:ch211-270 | atp1b1b | gsto1 | 4 |
| 134 | Ectoderm | Central Nervous System | Neuroblast | Forebrain | FBProg12 | FB | 1,2 | dmrt3a | eomesa | fezf2 | wnt8b | lhx5 | fezf1 | si:ch211-69b | rspo3 | zic6 | emx3 | cyp26c1 | wnt7bb | pdgfaa | tnfrsf19 | otx1 | si:ch211-285 | 2 |
| 135 | Unclear | Mesenchyme-related, organ muscle? | | | MesenUnk5 | MesenUnk5 | 5 | cxcl8b.3 | agrp | tmem98 | cxcl8b.1 | fgl2a | nt5e | colec11 | gcga | lurap1 | BX901920.1 | sfrp1b | si:ch211-106 | BX322618.1 | loxa | krt94 | zgc:110182 | 2 |
| 136 | Ectoderm | Central Nervous System | Neuron | Mid-hindbrain boundary, inhibitory | MHBNeurGABA12 | MHB | 1,2 | zfpm2b | nkx1.2lb | sox14 | slc6a1b | gad2 | tal1 | slc32a1 | lhx1a | gata2a | scrt1a | si:dkey-175g | gad1b | uncx | gata3 | lhx5 | celf4 | 1 |
| 137 | Ectoderm | Placode | | Lateral line | LatLine | LatLine | 1,2,5 | si:ch211-103 | fat1b | twf2b | si:ch211-172 | si:rp71-1c10. | mcf2lb | hmx2 | six2b | krt1-c5 | col17a1b | si:dkey-222f2 | hmx3a | si:dkeyp-110 | tagapa | si:dkey-16l2. | wnt2 | 6 |
| 138 | Ectoderm | Central Nervous System | Neuron | Retinal, differentiating neurons | RetDiff25f | RetDiff | 2,5 | sept4a | ndrg1b | CABZ010555 | cspg5b | vsx1 | crx | fkbp1ab | crabp1a | ptprua | si:dkey-238o | hes2.2 | foxn4 | foxg1b | otx5 | otx2b | sox12 | 1 |
| 139 | Ectoderm | Central Nervous System | Neuron, differentiating | Hindbrain | HBDiff1c | HB | 1 | jag1a | prdm12b | gbx1 | bhlhe22 | otpb | pcdh18a | ascl1b.1 | neurog1 | her13 | sox21a | sulf2a | sp8b | prdm8 | CR848812.1 | her8.2 | BX530077.1 | 0 |
| 140 | Mesoderm | Pharyngeal arch | | | Arch12 | Arch2 | 1,2 | tbx1 | kazald2 | gfra3 | pdgfab | meis3 | pdgfrb | tmem88b | prdm1a | meox1 | si:dkey-261h | osr1 | prrx1a | pcolcea | sfrp2 | foxc1a | efna5a | 1 |
| 141 | Ectoderm | Lens placode | Lens | | LensAll | Lens | 1,2,5 | BX548032.3 | nags | lim2.5 | si:ch211-254 | cryba1l2 | gja8b | crybb1l3 | cps1 | endou2 | cryba1a | si:ch73-111m | capn3a | lim2.3 | cx23 | crygn1 | tmprss5 | 2 |
| 142 | Ectoderm | Central Nervous System | Neuron, differentiating | Forebrain | FBDiffAll | FB | 1,2,5 | kctd4 | nkx2.4b | nkx2.1 | nkx2.4a | hmx2 | hmx3a | FP085414.1 | sox1a | ptpn5 | scg2b | gpr85 | six6a | nexmifb | sox14 | sox1b | celf4 | 0 |
| 143 | Ectoderm | Central Nervous System | Neuron | Spinal cord motor neurons | SCNeurMN12b | SC | 1,2 | chata | slc5a7a | mnx1 | slc18a3a | si:ch211-247 | chodl | chrnb4 | tac1 | lhx4 | si:ch211-113 | tmie | ssx2ipb | l1cama | gfra3 | ret | syn1 | 2 |
| 144 | Mesoderm | Kidney | Convoluted tubule? | | Kidney | Kidney | 5 | slc9a3.1 | AL929208.1 | zgc:195356 | amn | si:dkey-194e | ggt1b | cldn10b | si:ch211-274 | mfsd4ab | cubn | c1qtnf9 | slc5a9 | hnf1bb | pdzk1ip1 | si:dkey-39a1 | prr15lb | 4 |
| 145 | Ectoderm | Integument | Epidermis | Fin basal cells | BasalFin12 | BasalEpiFin1 | 1,2 | itih5 | bhlha9 | tspearb | adamtsl7 | srpx2 | sp6 | glsl | vcana | msx2a | hapln3 | hhipl2 | and3 | mxc | dkk1a | col4a3 | dlx2b | 0 |
| 146 | Ectoderm | Integument | Integument Gill? | Gill | Gill5c | Gill | 5 | CU062645.1 | si:dkey-56e3. | si:dkey-239b | si:dkey-7c18. | si:ch211-226 | CABZ010489 | si:dkey-239b | tkta | AL928650.1 | vwa10.1 | psmb8a | caspbl | jac10 | atp10b | tmem176l.3 | si:dkey-228d | 6 |
| 147 | Mesoderm | Lateral plate mesoderm | Heart primordium | | HeartPrimord | Heart | 1 | CR762480.2 | si:ch73-190f | FO704871.1 | piwil2 | astn1 | zgc:171482 | clec19a | gata5 | pitx1 | sema3e | emilin2b | osr2 | col6a4a | osr1 | si:ch211-160 | BX001014.2 | 3 |
| 148 | Ectoderm | Neural crest | Cranial neural crest | Cartilage | NCcart5b | NCcart5 | 5 | itih1 | ninj2 | CLEC3A | pcolceb | wisp3 | acana | mia | matn1 | snorc | tnxba | arsia | loxl4 | si:dkey-6n6.1 | chad | ptgdsa | si:dkey-31g6. | 2 |
| 149 | Mesoderm | Notochord | | | Notochord25 | Notochord | 2,5 | si:dkey-234i1 | si:ch211-13f | matn3b | paqr4b | chadla | igsf10 | slc34a1a | zgc:175280 | susd5 | si:dkey-31g6. | tbx3b | LO018422.1 | gas2b | tbxta | si:dkey-6n6.1 | chad | 4 |
| 150 | Mesoderm | Myeloid lineage | Leukocyte | Neutrophil | Neutrophil | Neutrophil | 1,2,5 | si:dkey-42l23 | c6ast1 | si:dkey-238m | itln1 | CR354540.1 | pigrl2.3.1 | il4 | si:ch211-8c1 | ch25hl2 | si:ch73-170d | ANPEP (1 of | si:dkey-53k1 | ctss1 | glipr1a | dennd3b | si:dkey-19b2 | 5 |
| 151 | Ectoderm | Central Nervous System | Neuron, differentiating | Mid-hindbrain boundary | MHBDiff1 | MHB | 1 | helt | zgc:101100 | gsx1 | otx2a | gata2a | BX530077.1 | tfap2e | ascl1b.1 | inavaa | gadd45gb.1 | nkx2.2a | her13 | otx1 | CR751602.2 | ascl1a | nes | 1 |
| 152 | Endoderm | Intestine | Intesinal epithelial cell | Intestinal bulb, enterocyte? | IntestineEpi5a | IntestineEpi | 5 | enpp7.1 | si:ch211-142 | slc28a1 | sult1st2 | cpo | alpi.2 | rh50 | mogat2 | si:ch211-113 | adgrg4a | apoa4b.3 | slc34a2a | zgc:136472 | zgc:154142 | ace2 | chs1 | 4 |
| 153 | Mesoderm | Blood | Hematopoietic cell | Progenitor | RBCProgAll | RBCprog | 1,2,5 | fermt3b | top1 | s1pr5b | tagapb | si:ch211-103 | TCIM (1 of m | drl | zar1 | hif1al2 | si:dkey-261j4 | lrrc15 | si:dkey-93h2 | si:dkey-261j4 | si:cabz01074 | cldng | klf1 | 5 |
| 154 | Mesoderm | Muscle | Cephalic muscle | | MusCeph | MusCeph | 5 | zgc:66156.1 | fhl5 | FO704758.1 | zgc:172270 | tnni2a.2 | trim54 | sync | mybpc2a | myhb | tnni2a.3 | smpdl3b | myhz1.1.1 | mybpc1 | mybpha | zgc:136908 | pdlim3a | 3 |
| 155 | Mesoderm? | Tailbud | | | Tailbud | Tailbud | 1 | si:ch211-196 | si:dkey-14o1 | hoxb13a | hoxc13b | hoxc12b | hoxa13a | hoxc13a | sp5l | efemp2b | hoxd13a | hoxa13b | BX005254.4 | angptl2a | pcdh8 | c1qtnf5 | msgn1 | 2 |
| 156 | Ectoderm | Neural crest | Xanthophore1 | | NCxantho1 | NCxantho | 1 | sprb | zgc:113337 | slc2a15a | zgc:113142 | mlphb | bscl2l | pts | plin6 | cax1 | si:dkey-251i1 | cyb561a3b | aox5 | slc22a7a | tmem130 | slc2a15b | ednrba | 3 |
| 157 | Mesoderm | Blood | Hematopoietic cell | Nucleate erythrocyte | RBC5 | RBC5 | 5 | si:ch211-103 | si:ch211-250 | nt5e | slc6a22.2 | si:ch211-207 | alas2 | hif1al2 | zgc:163057 | MFAP4 (1 of | CR293511.1 | epb41b | hemgn | nmt1b | nt5c2l1 | hbae5 | klf1 | 4 |
| 158 | Mesoderm | Notochord | | | Notochord1 | Notochord | 1 | wu:fc23c09 | susd5 | si:ch211-13f | paqr4b | chadla | si:dkeyp-118 | LO018422.1 | tbx3b | matn3b | igsf10 | si:dkey-6n6.1 | tbxta | loxl5b | gas2b | si:dkey-234i1 | adra2da | 4 |
| 159 | Nonspecific genes | | | | NonspecificAll | Nonspecific | 1,2,5 | NC_002333.1 | pax6b | mt-nd5 | fabp11a | aifm1 | CT027638.1 | cad | zgc:158463 | pola1 | cox17 | si:ch211-282 | ddx24 | CU467861.1 | mt-nd3 | ddx21 | pclaf | 2 |
| 160 | Ectoderm | Central Nervous System | Neuron, differentiating | Spinal cord | SCDiff12c | SC | 1,2 | pkd1b | cdx4 | exd1 | grm2b | fhdc1 | jag1a | si:dkeyp-110 | pou3f2a | ldlrad2 | prdm12b | dbx1b | lrrc4.1 | lbx1b | si:ch211-170 | mb21d2 | hoxc9a | 2 |
| 161 | Ectoderm | Neural crest | Cranial neural crest | | NCcranial1d | NCcran1 | 1 | npb | ednrab | sema3e | cxcr4a | arhgap36 | ccdc80l2 | jcada | cxcl12b | prrx1b | CU570769.1 | csf1a | grem2b | twist3 | inka1a | ednraa | fli1a | 0 |
| 162 | Mesoderm | Spleen | Spleen mesoderm | | Spleen | Spleen | 5 | lepb | BX511186.1 | si:ch1073-11 | irak3 | tnfaip2b | saa | si:ch211-198 | cxcl8a | steap4 | snap23.2 | si:ch211-132 | ms4a17a.10 | BX908750.1 | tlr5b | ch25h | hamp | 3 |
| 163 | Ectoderm | Central Nervous System | Neuroblast | Roofplate?, hindbrain | Roofplate | Roofplate | 1,2,5 | lmx1a | si:dkey-1c11. | BX469902.1 | hepacama | fabp7b | zgc:114181 | bcl6b | AL928901.1 | bmp2a | lmx1ba | gdpd5a | ppfibp2a | cfap45 | rspo2 | si:ch211-66e | cfap99 | 3 |
| 164 | Ectoderm | Neural crest | Cranial neural crest | | NCcranial1e | NCcran1 | 1 | irx6a | si:dkey-250k1 | ednrab | crestin | artnb | mmp17b | fgf10a | prdm1a | pdgfbb | gfra3 | gpc1b | apela | phlda3 | prrx1b | grem2b | meis3 | 1 |
| 165 | Ectoderm | Integument | Integument | Ionocyte? | IonoNCC5b | IonoNCC | 5 | si:ch211-180 | zgc:165423 | calhm1 | si:dkeyp-75b | pkhd1l1 | nox1 | CT030129.1 | si:ch1073-15 | si:ch73-329n | hhla2b.1 | ca15c | mslna | krt1-19d | MFAP4 (1 of | ca15a | cyldb | 5 |
| 166 | Ectoderm | Central Nervous System | Neuroblast | Anterior spinal cord | antSCProg1c | antSC | 1 | hoxb6b | hoxb5b | hoxb6a | dhrs3b | hoxc1a | hoxb1b | prdm8 | hoxb5a | ncam1b | si:ch73-265d | sulf2a | hoxd4a | foxb1a | foxb1b | sox21a | rxrga | 1 |
| 167 | Mesoderm | Lateral plate mesoderm | Vasculature | Vessel | Vessel2 | Vessel2 | 2 | si:dkeyp-106 | zgc:158423 | fpr1 | FO704741.1 | apof | lamc3 | cldn11a | zgc:162612 | ndnfl | tgm2l | chst2b | si:ch211-160 | calcrla | CABZ010732 | osr1 | pdgfrl | 4 |
| 168 | Endoderm | Intestine | Intesinal epithelial cell | Mid? | IntestineEpi5b | IntestineEpi | 5 | tspan34 | si:ch211-202 | plac8.2 | itga6l | si:dkey-188i1 | zgc:77748 | cdx1b | ca4b | ccl25a | zgc:136872 | zgc:158432 | anxa2b | clrn3 | zgc:198329 | si:ch73-194h | cldn15la | 6 |
| 169 | Ectoderm | Integument | Integument | Mucosal? | MucNCC | MucNCC | 1,2,5 | BX649522.1 | st6galnac1.1 | zgc:112433 | muc5.2 | muc5.1 | si:dkey-65b1 | slc46a3 | si:ch211-163 | fev | si:dkey-246j7 | pou2f3 | p2rx1 | s1pr5b | st6gal2b | C1GALT1 (1 c | fam83fa | 4 |
| 170 | Ectoderm | Neural crest | Cranial neural crest | | NCcranial5c | NCcran5 | 5 | cald1b | si:ch211-286 | rgs5a | rgs4 | twist2 | frzb | postna | asip2b | tat | vcanb | si:dkey-12l12 | pah | aspn | si:ch211-106 | hpdb | ttc36 | 3 |
| 171 | Mesoderm | Visceral mesoderm | Vascular smooth muscle | | MusVascSmo | MusVascSmo | 5 | pdlim3a | csrp1b | desmb | mylkb | gucy1a1 | cabp1a | CR848749.2 | fhl1a | cald1b | CU571069.1 | smtna | zgc:110699 | myh11a | BX908750.1 | kcne4 | pde9a | 1 |
| 172 | Ectoderm | Central Nervous System | Neuron, differentiating | Hindbrain | HBDiff2 | HB | 2 | sox14 | cyp2aa2 | gata2a | vsx1 | hes2.2 | foxn4 | si:ch211-117 | cited1 | lhx1a | ptprua | gdf11 | cdkn2c | CR848844.1 | bhlhe22 | golga7ba | foxp2 | 1 |
| 173 | Ectoderm | Neural crest | Neuronal/Glial | | NCneural25 | NCneural | 2,5 | si:ch211-197 | zwi | scn4ab | hepacama | dkk1a | nr4a2b | hbegfb | sox10 | zgc:112437 | mcamb | adgrg6 | foxd3 | ccdc80 | pcdh7a | ngfrb | tp53i11b | 2 |
| 174 | Ectoderm | Neural crest | Melanophore1 | | NCmel1 | NCmel | 1 | opn5 | tyr | slc24a5 | tspan10 | si:dkey-24c2. | ednrba | crestin | cdk15 | aqp7 | slco1c1 | zgc:91968 | pts | rab34b | tmem125b | cax1 | emp3b | 2 |
| 175 | Unclear | Placode-related, epithelial, gor | Pronephros epi? | | PlacodalUnk | PlacUnk | 1 | si:ch211-256 | zgc:112408 | vill | gfral | CABZ010730 | gbgt1l3 | si:ch211-130 | st3gal1l2 | gng8 | zgc:101731 | CR759836.1 | gcnt7 | ddx4 | tph1b | prr15la | cx28.8 | 4 |
| 176 | Ectoderm | Central Nervous System | Floor plate | Spinal cord | FloorplateSCAll | Floorplate | 1,2,5 | wnt4b | urp2 | vwa3a | spon1a | gucy1a1 | rnf32 | s100b | cdkl1 | cfap157 | tbata | slit1b | si:ch211-215 | npr3 | morn3 | shhb | ccdc169 | 0 |
| 177 | Ectoderm | Central Nervous System | Neuron | Retinal, mix of neuron types? | RetNeuron5 | RetNeuron | 5 | si:ch211-13f | lrit1a | nyx | gabra4 | gabrr2a | RIC3 | cabp5a | igsf21b | si:ch211-232 | nrn1lb | mfsd2b | rgs11 | FAM163A | lactbl1b | trpm1a | si:ch211-160 | 3 |
| 178 | Ectoderm | Neural crest | Melanophore2 | | NCmel25 | NCmel | 2,5 | slc30a8 | slc38a11 | alkal2b | si:dkey-24c2. | zdhhc2 | CLEC3A | slc24a5 | tyr | zgc:91968 | slc24a4a | si:dkeyp-118 | prkar1b | tspan10 | rgs16 | fmn2a | emp3b | 3 |
| 179 | Endoderm | Pharyngeal endoderm | Pharyngeal epidermis | | PharEpi2 | PharEpi2 | 2 | shhb | nkx3.3 | pitx1 | sp6 | shha | foxa2 | ociad2 | inhbaa | abi3b | vgll4l | pitx2 | nkx2.3 | afap1l1a | frem2a | tp63 | CABZ010244 | 0 |
| 180 | Endoderm | Pancreas | Pancreatic epsilon cell | | PancExo | PancExo | 5 | CELA1 (1 of n | si:dkey-266f7 | si:ch73-44m | pglyrp6 | BX908782.1 | CELA1 (1 of r | CELA1 (1 of n | si:ch211-255 | endou | CELA1 (1 of r | c6ast3 | sycn.3 | dnase1 | CELA1 (1 of n | si:ch211-240 | syt15 | 4 |
| 181 | Ectoderm | Neural crest | Cranial neural crest | | NCcranial5d | NCcran5 | 5 | pax1b | fgfbp2a | pax1a | abtb2a | mctp1a | dlx4a | fbln7 | sost | cyp2ad2 | ecrg4a | nog3 | dlx1a | bmp2a | gata2a | dlx6a | rgs5b | 0 |
| 182 | Ectoderm | Integument | Periderm | Sensory? | Peri2c | Peri2 | 2 | si:dkey-56i24 | si:ch211-264 | si:dkey-239b | ponzr4 | zgc:163121 | wu:fj05g07 | si:dkey-9l20. | muc13a | si:ch211-264 | irg1l | si:ch211-153 | ponzr3 | ano9a | jac10 | grk5 | zgc:123217 | 8 |
| 183 | Ectoderm | Integument | Periderm | Sensory? | Peri1b | Peri1 | 1 | si:ch1073-15 | BX511250.1 | BX936456.1 | si:ch211-153 | si:dkeyp-67a | LO018208.1 | irg1l | he1.3 | cyp2aa8 | prss60.2 | si:ch73-181m | slc35a3b | ccl20a.3 | tex36 | cyp2aa11 | si:dkey-160o | 5 |
| 184 | Mesoderm | Myeloid lineage | Monocyte | Macrophage | Macro12 | Macro | 1,2 | si:dkey-32n7. | cx32.2 | CABZ010414 | si:ch73-343l4 | entpd2a.1 | gpr84 | slc2a6 | plaua | si:ch73-343l4 | fcer1g | CABZ010800 | si:dkey-27h1 | dicp3.1 | cd22 | MFAP4 (1 of | CABZ010881 | 4 |
| 185 | Ectoderm | Neural crest | Xanthophore | | NCxantho25 | NCxantho | 2,5 | oacyl | zgc:113142 | cyb561a3b | bscl2l | plin6 | si:dkey-251i1 | cax1 | slc22a7a | tmem130 | aox5 | slc2a15a | si:dkey-126g | slc2a11b | slc2a15b | mlpha | pts | 3 |
| 186 | Ectoderm | Integument | Epidermis | Basal cells | Basal5b | BasalEpi5 | 5 | zgc:136254 | adamtsl7 | zgc:173443 | si:ch211-167 | wnt5a | bhlha9 | si:ch211-182 | mslna | BX004816.1 | si:ch211-251 | and3 | si:ch73-181m | si:dkey-56e3. | sp6 | zgc:123217 | NPNT (1 of m | 8 |
| 187 | Nonspecific genes | | | | Nonspecific1a | Nonspecific | 1 | six3b | fabp11a | fahd1 | znhit3 | zgc:194224 | thumpd3 | umps | ppat | dctpp1 | rrp8 | noc3l | qtrt2 | cmss1 | myca | exosc8 | polr1e | 1 |
| 188 | Ectoderm | Integument | Epidermis | Fin basal cells | BasalFin5b | BasalEpiFin5 | 5 | hhipl2 | si:ch211-76l2 | sod3b | col17a1a | plek2 | slc6a11a | cxcl18a.1 | si:dkey-95h1 | si:dkey-119m | mmp30 | tkta | cxl34b.11 | tnfsf10l4 | bgnb | entpd3 | CABZ010681 | 3 |
| 189 | Endoderm | Pancreas | Pancreatic B, Pancreatic A | | PancAB | PancAB | 1,2,5 | gck | scgn | a1cf | adm2a | rprma | c2cd4a | renbp | pdx1 | gcgb | fev | npas4a | si:ch211-163 | si:ch211-113 | emb | tmem63c | ndufa4l2a | 2 |
| 190 | Ectoderm | Integument | Periderm | Mucosal? | Peri1c | Peri1 | 1 | muc13a | si:dkey-56i24 | si:dkey-239b | si:ch211-153 | tex36 | irg1l | zgc:123217 | mia | trim25l | si:dkey-9l20. | si:ch211-153 | krt1-19d | si:ch211-264 | rab11al | si:dkey-160o | zgc:113442 | 9 |
| 191 | Mesoderm | Paraxial mesoderm | Parcordal cartilage | | Parachord1 | Parachord | 1 | ndufa4l2a | itga11a | nav1b | cd81b | apof | foxd5 | vstm4b | has2 | CABZ010732 | ndnfl | foxl1 | vcana | nkx3.2 | si:dkeyp-106 | pcolce2a | zgc:162612 | 2 |
| 192 | Ectoderm | Central Nervous System | Neuron, differentiating | Forebrain | FBDiff12 | FB | 1,2 | sim1a | l1cama | fezf1 | FP085414.1 | lhx5 | otpb | uncx | fezf2 | slc17a6b | celf4 | otpa | nrn1a | adcyap1b | nell2b | adam22 | mpp2a | 0 |
| 193 | Nonspecific genes | | | | Nonspecific1b | Nonspecific | 1 | umps | six3b | exosc8 | ftsj3 | cmss1 | FP245462.1 | impdh1b | dctpp1 | wdr12 | myca | mtbl | ppan | paics | pold2 | pus7 | chtf8 | 0 |
| 194 | Ectoderm | Central Nervous System | Neuron, differentiating | Spinal cord, floor plate? | SCDiff12c | SC | 1,2 | wnt4b | spon1a | pth1a | cyr61l2 | tbata | cfap126 | spata18 | nkx2.2b | si:ch211-176 | cfap57 | cfap299 | armc4 | CABZ010720 | cfap58 | nkx2.9 | zgc:113531 | 2 |
| 195 | Ectoderm | Central Nervous System | Neuroblast | Mid-hindbrain boundary | MHBProg1 | MHB | 1 | pax5 | en2a | en2b | en1b | sox21a | pax2a | cnpy1 | qdpra | nanos1 | exosc8 | dctpp1 | dnajc2 | her8a | sox19a | pou3f3a | hpf1 | 0 |
| 196 | Ectoderm | Integument | Taste bud | | TasteBud | TasteBud | 5 | si:dkeyp-92c | rgs1 | BX005256.1 | zgc:194948 | PLCB2 | CABZ010684 | muc5.3 | muc5f | gng13a | trpm5 | avil | ceacam1 | LO018550.2 | kcnk17 | zgc:158412 | prss60.2 | 3 |
| 197 | Mesoderm | Visceral mesoderm | Enteric smooth muscle | | MusEntSmo | MusEntSmo | 5 | kcne4 | mfap5 | myh11a | mylka | foxf1 | mylkb | bgnb | si:ch211-270 | acta2 | foxl1 | loxa | BX908750.1 | lmod1b | tagln | BX088707.3 | CU914776.1 | 1 |
| 198 | Ectoderm | Placode | Neuron | Hair cell | HairCell | HairCell | 1,2,5 | tmc2a | dnajc5b | si:ch211-229 | si:dkey-33m1 | si:ch211-284 | si:ch73-261i2 | CR792457.1 | tmc2b | atp6v1c2 | zmp:0000001 | lhfpl5b | baiap2l2b | tspan1 | sh2d4ba | zgc:171927 | si:dkey-103g | 5 |
| 199 | Ectoderm | Placode | Adenohypophyseal | Hypophysis | Hypophysis | Hypophysis | 5 | tmtops2a | gbe1a | lrrc58a | ccka | fgl2a | FO681288.1 | ptger2b | tubb6 | pmch | tkta | pitx1 | rerglb | bco2a | tnmd | lmx1ba | bco2l | 0 |
| 200 | Ectoderm | Lens placode | Lens | | Lens12 | Lens | 1,2 | crygm2d7 | crygmxl2 | crygm2d17.1 | crygm2d10 | crybb1l1 | crygm2d8.1 | crybb1l2 | crygm2d13 | cryba1l1 | cryba1b | crybb1 | crygn2 | cryba2a | cryba2b | cryba4 | fabp11a | 0 |

| | | | | | | | | | | | | | | | | | | | | | | | | |
|---|---|---|---|---|---|---|---|---|---|---|---|---|---|---|---|---|---|---|---|---|---|---|---|---|
| 201 | Mesoderm | Lateral plate mesoderm | Vasculature | Vessel | Vessel5 | Vessel5 | 5 | zmp:0000001 | slc13a4 | slc22a7b.1 | si:ch211-76l2 | zgc:158423 | si:dkeyp-106 | slc47a1 | slc13a2 | plp1b | slc47a2.1 | sat1a.1 | htr2cl1 | clec3ba | CABZ010732 | apof | cnnm2b | 3 |
| 202 | Ectoderm | Central Nervous System | Neuron | Ciliated, spinal cord | SCNeurCilia12b | SC | 1,2 | **pkd2l1** | **pkd1l2a** | si:ch1073-70 | rnf220b | grm2a | c2cd4a | syt6b | fam19a1a | **urp1** | **urp2** | flj13639 | si:ch211-258 | kcnf1a | sst1.1 | dcdc2b | nppc | 2 |
| 203 | Mesoderm | Pharyngeal arch | | | Arch2b | Arch2 | 2 | **lhx6** | **gsc** | foxl1 | hoxb6a | **barx1** | **foxf2a** | **hoxb5b** | **hoxb5a** | CABZ010751 | tmeff2a | **prrx1b** | rspo2 | **dlx1a** | **meis3** | aldh1a2 | **vasnb** | 0 |
| 204 | Mesoderm | Pectoral fin bud | Progenitor? | | FinBud2 | FinBud | 2 | **glsl** | **angptl6** | eve1 | hoxb13a | **ptx3a** | **hoxa13a** | si:ch211-14k | c1qtnf5 | hapln1b | hoxc13a | oscp1a.1 | adgrg6 | **hoxa13b** | hoxc13b | **mxra8a** | **loxa** | 1 |
| 205 | Mesoderm | Lateral plate mesoderm | Heart primordium | Cardiac muscle | MusCard | MusCard | 1,2 | **lft2** | **cx36.7** | **nppa** | **slc8a1a** | **nppb** | **adprhl1** | **ryr2b** | **tnnc1a** | **myh6** | lmod2b | **myh7l** | **lrrc10** | pln | **csrp3** | **nkx2.5** | **myh7** | 0 |
| 206 | Ectoderm | Integument | Epidermis | Basal cells | Basal2b | BasalEpi2 | 2 | BX855590.1 | zgc:100997 | ponzr4 | cdh26.2 | **mxc** | CU459012.1 | si:ch211-153 | si:dkey-95h1 | ubap1la | cers3a | col28a1a | si:ch211-241 | **col17a1b** | si:ch211-264 | grk5 | jac10 | 5 |
| 207 | Mesoderm | Hypoblast | Hatching gland | | HG2 | Hatching | 1 | BX000534.1 | BX322603.1 | vtcn1 | zmp:000000 | si:dkey-26g8. | zp3.2.1 | CU695215.2 | slc30a8 | fmn1 | LO017965.1 | cx30.9 | CT033796.2 | si:ch211-171 | **mctp2b** | sytl4 | zgc:101785 | 3 |
| 208 | Ectoderm | Integument | Integument Ionocyte | vH Ionocyte | IonovH | IonovH | 1,2,5 | si:dkey-192d | ceacam1 | drd4-rs | **gcm2** | etf1a | **foxi3a** | **slc4a1b** | prg4a | **ca15a** | agrp | si:ch211-182 | cldn17 | zgc:193726 | si:ch73-359m | si:ch73-14h1. | scgn | 5 |
| 209 | Mesoderm | Muscle | Skeletal muscle | Slow muscle cell | MusSkelSlow1b | MusSkelSlow | 1 | si:dkey-250i3 | mylk4a | **fhl1a** | smyhc3 | klhl41a | **foxd5** | CABZ010817 | **smyhc2** | CU633479.2 | alpk3b | **flncb** | **chrng** | mmrn2b | BX649294.1 | CR381686.5 | AL772146.2 | 1 |
| 210 | Mesoderm | Spleen | Spleen epithelial cells? | | SpleenEpi | SpleenEpi | 1 | ggt1a | **tnfa** | mmp13a | lacc1 | **lepb** | sult5a1 | plaua | cd44a | si:dkey-19b2 | **csf3b** | nox1 | mmp13a.1 | si:ch211-39f | ms4a17a.4 | scpp8 | si:ch1073-11 | 3 |
| 211 | Ectoderm | Pineal gland | Epiphysis | | Epiphysis | Epiphysis | 1 | rdh20 | cidea | plin1 | **rrh** | CR792417.1 | slc16a8 | tmem72 | selenou1b | slc4a5 | BX323820.1 | **trpm1a** | inpp5kb | slc13a4 | msnb | slc22a6l | cracr2ab | 0 |
| 212 | Mesoderm | Myeloid lineage | Monocyte | Macrophage | Macro1 | Macro | 1 | si:dkey-1c7.2 | si:ch211-225 | xcr1a.1 | FP085399.5 | gpr34b | tlr1 | si:ch211-11c | si:dkey-27j5. | si:dkey-83f18 | CR626902.1 | spic | si:ch73-343g | il1fma | zgc:123107 | FP085399.2 | si:ch211-156 | 8 |
| 213 | Mesoderm | Blood vessel | Endothelia | | Endothe1 | Endothelial | 1 | f8 | vwf | si:ch211-33e | ccm2l | ecscr | **flt1** | **tie1** | sele | **clec14a** | **gpr182** | myct1b | gata5 | myct1a | edn2 | pecam1 | si:ch211-145 | 2 |
| 214 | Endoderm | Intestine | Intesinal epithelial cell | | IntestineEpi1 | IntestineEpi | 1 | **chia.3** | si:ch211-77g | wu:fa56d06 | ACOT12 | zgc:153968 | si:dkey-236e | **mogat2** | apoea | **slc34a2a** | si:ch211-142 | slc6a19b | slc13a2 | si:dkey-103j1 | pdzk1 | zgc:158846 | **aqp8a.2** | 6 |
| 215 | Ectoderm | Integument | Periderm | | Peri1d | Peri1 | 1 | si:dkey-39a1 | si:dkeyp-67a | trim25l | abca4a | zgc:113442 | **grhl3** | si:ch211-264 | chrna5 | dpys | insl5a | zbtb7a | ponzr5 | BX323590.2 | LO017816.1 | **noxo1a** | si:dkey-222f2 | 5 |
| 216 | Mesoderm | Connective tissue | Osteoblast | | Osteo | Osteo | 2,5 | **bgna** | **entpd5a** | si:ch211-106 | col5a2b | sgms2a | **panx3** | ifitm5 | si:dkey-32e6. | si:ch211-170 | **spp1** | enam | bmp8a | zgc:92162 | tgfb2l | chst3b | slc8a4b | 3 |
| 217 | Endoderm | Liver | Hepatocyte?? | Kupffer cells? | Hepato5c | Liver | 5 | si:dkey-79f11 | BX927210.1 | CABZ010006 | si:ch211-113 | si:dkey-7f3.1 | FO834888.1 | CU856343.1 | gltpd2 | rln3a | acod1 | **soat2** | apoc4 | **gpr84** | **hp** | si:ch211-271 | **c3b.1** | 4 |
| 218 | Endoderm | Hypochord | | | Hypochord | Hypochord | 1 | si:ch211-157 | **spon2a** | pappaa | pgfb | col8a1b | **mnx1** | **ptx3a** | **matn3b** | **npr3** | cyr61l2 | **angpt1** | loxl5b | **cd59** | aplnr2 | ism2a | sim1a | 1 |
| 219 | Germ | Germ cell | Primordial germ cell | | PGC | PGC | 1 | **tdrd7a** | **dazl** | fkbp6 | zpcx | tdrd12 | **dnd1** | **ca15b** | pld6 | adad1 | **cpeb1b** | **nanos3** | akap1a | si:dkey-189g | CR382383.2 | zgc:112416 | si:ch211-226 | 3 |
| 220 | singleton | | | | Nonspecific | Nonspecific | | | | | | | | | | | | | | | | | | |
| | | | | | | | | | | | | | | | | | | | | | AVG | 2.2 |
| | | | | | | | | | | | | | | | | | | | | | STDEV | 2.1 |
